# Colonization with the commensal fungus *Candida albicans* perturbs the gut-brain axis through dysregulation of endocannabinoid signaling

**DOI:** 10.1101/2020.02.27.968198

**Authors:** Laura Markey, Andrew Hooper, Laverne C. Melon, Samantha Baglot, Matthew N. Hill, Jamie Maguire, Carol A. Kumamoto

## Abstract

Anxiety disorders are the most prevalent mental health disorder worldwide, with a lifetime prevalence of 5-7% of the human population. Although the etiology of anxiety disorders is incompletely understood, one aspect of host health that affects anxiety disorders is the gut-brain axis. We used a mouse model of gastrointestinal (GI) colonization to demonstrate that the commensal fungus *Candida albicans* affects host health via the gut-brain axis. In mice, bacterial members of the gut microbiota can influence the host gut-brain axis, affecting anxiety-like behavior and the hypothalamus-pituitary-adrenal (HPA) axis which produces the stress hormone corticosterone (CORT). Here we showed that mice colonized with *C. albicans* demonstrated increased anxiety-like behavior and increased basal production of CORT as well as dysregulation of CORT production following acute stress. The HPA axis and anxiety-like behavior are negatively regulated by the endocannabinoid anandamide (AEA). *C. albicans-colonized* mice exhibited systemic changes in the endocannabinoidome, within the GI tract and the brain, and showed a negative correlation between brain AEA levels and serum CORT. Further, increasing AEA levels using the well-characterized fatty acid amide hydrolase (FAAH) inhibitor URB597 was sufficient to reverse both neuroendocrine phenotypes in *C. albicans-colonized* mice. Thus, a commensal fungus that is a common colonizer of humans had widespread effects on the physiology of its host. To our knowledge, this is the first report of microbial manipulation of the endocannabinoid (eCB) system that resulted in neuroendocrine changes contributing to anxiety-like behavior.

## 1. Introduction

Anxiety disorders are the most prevalent mental health disorder worldwide, with a lifetime prevalence of 5-7% of the human population (Baxter et al., 2013). The etiology of anxiety disorders is complex and incompletely understood, although many different genetic and environmental factors have been identified (Leonardo and Hen, 2008). The gut-brain axis, the bi-directional communication between the gut and the brain through multiple host systems, has been shown to affect anxiety disorders (Luna and Foster, 2015). Constant communication between the periphery and the brain is required for normal homeostasis and overall health.

One component of the gut-brain axis thought to play a role in anxiety disorders is the hypothalamus-pituitary-adrenal (HPA) axis. The endocrine output of the HPA axis, the stress hormone corticosterone (CORT), is a broad regulator of host health including the immune, neuroendocrine, metabolic and cardiovascular systems. Under basal conditions, CORT release is regulated by the circadian rhythm and helps to maintain homeostasis (Dallman et al., 1993). CORT is also a key part of the acute psychological stress response and a rapid increase in circulating CORT is essential for mounting an effective response to stress (Sapolsky et al., 2000). Dysregulation of the HPA axis and CORT production is correlated with mental health disorders including anxiety and depression (Kallen et al., 2008).

The gut microbiota, the diverse assemblage of microbes that colonizes the GI tract of humans, has been shown to affect virtually every aspect of human health, including the gut-brain axis. Researchers using germ-free mouse models demonstrated that the gut microbiota plays an important role in the development and regulation of the HPA axis (Sudo et al., 2004). Studies have also shown that specific probiotic bacterial species can affect the gut-brain axis, as treatment with *Bifidobacterium sp*. and *Lactobacillus sp*. decreases anxiety-like behavior and normalizes HPA axis function in mice (Messaoudi et al., 2011). Investigation into the contribution of the gut-microbiota-brain axis to mental health could reveal novel regulators of anxiety-like behavior and offer mechanistic insight into the etiology of the disease.

*Candida albicans* is a commensal fungus that colonizes the GI tract of ~60% of the human population (Raimondi et al., 2019). Although a major colonizer of humans, its role as a commensal microbe is largely uncharacterized. Previous researchers have shown that GI *C. albicans* colonization induces local and systemic immune changes which are protective against systemic *C. albicans* infection and disease (Shao et al., 2019), and against infection with bacterial pathogens such as *C. difficile* (Markey et al., 2018) and *S. aureus* (Shao et al., 2019). The research that follows demonstrates that *C. albicans* affects host health beyond the GI tract. We show that *C. albicans* dysregulated HPA axis activity and increased anxiety-like behavior in a mouse model by altering endocannabinoid signaling.

The mammalian endocannabinoid (eCB) system regulates both the HPA axis and anxiety-like behavior. It consists of two major neuroactive lipids, N-arachidonoylethanolamide (AEA) and 2-arachidonoylglycerol (2-AG) and their receptors CB1 (distributed throughout the nervous system) and CB2 (present primarily on immune cells) (Devane et al., 1992; Mechoulam et al., 1995). AEA and 2-AG are produced in response to activation in post-synaptic neurons, and act at CB1 on pre-synaptic neurons to limit neurotransmitter release. AEA and 2-AG levels are regulated by substrate availability and the activity of their synthetic and degradative enzymes (Placzek et al., 2008). Using genetic and pharmacological tools in rodents, researchers have demonstrated that manipulation of CB1 signaling affects basal and stress-induced activation of the HPA axis as well as performance in tests for anxiety-like behavior (Barna et al., 2004; Hill et al., 2011). Here we show that treatment with the FAAH inhibitor URB597, which increases AEA levels and thus CB1 signaling, is sufficient to reverse the effect of *C. albicans* colonization on CORT and anxiety-like behavior. We also show that colonization with *C. albicans* altered levels of eCBs in the gut and that brain AEA levels correlated with the increase in serum CORT observed in the *C. albicans*-colonized mice. These results establish a mechanism by which a specific member of the gut microbiota, *C. albicans*, affects activity in the brain.

## 2. Results

### 2.1 A single oral inoculation with *Candida albicans* is sufficient to establish gastrointestinal colonization without inflammatory disease

We used a murine colonization model to investigate the effects of *C. albicans* inoculation on the mammalian host. Co-housing of up to 23 5-week-old mice in a large cage was used to standardize gut microbiota by coprophagy and therefore only female mice were used in these studies. The mice were acclimated to our facility and handled daily while co-housed. They were then either inoculated with a single dose of *C. albicans* or mock-inoculated with buffer and transferred to small cages in groups of 3-4. *C. albicans* was measurable in fecal pellets after 24h and 3 days after inoculation (Fig. 1B) and was detected throughout the GI tract (Fig. S1). We assessed behavior two days post-inoculation and mice were sacrificed after three days of colonization (Fig. 1A). *C. albicans-colonized* mice did not exhibit symptoms of illness or lose >5% bodyweight (Fig. S2). A multiplex ELISA was used to measure inflammatory cytokines in the serum after sacrifice and low levels of circulating IL-1β, IFN-γ, IL-10 and IL-6 were measured in mock-inoculated and *C. albicans-colonized* mice (Fig. 1C). Although IL-6 was significantly increased in *C. albicans-colonized* mice (Student's t-test, p=0.041), IL-6 as well as all other cytokines were well below the levels typically observed in mice with *C. albicans* disease (Tuite et al., 2005). Thus, our model resulted in stable colonization of mice with *C. albicans* in the GI tract that was not associated with disease, modeling commensal GI colonization.

**Figure 1:**
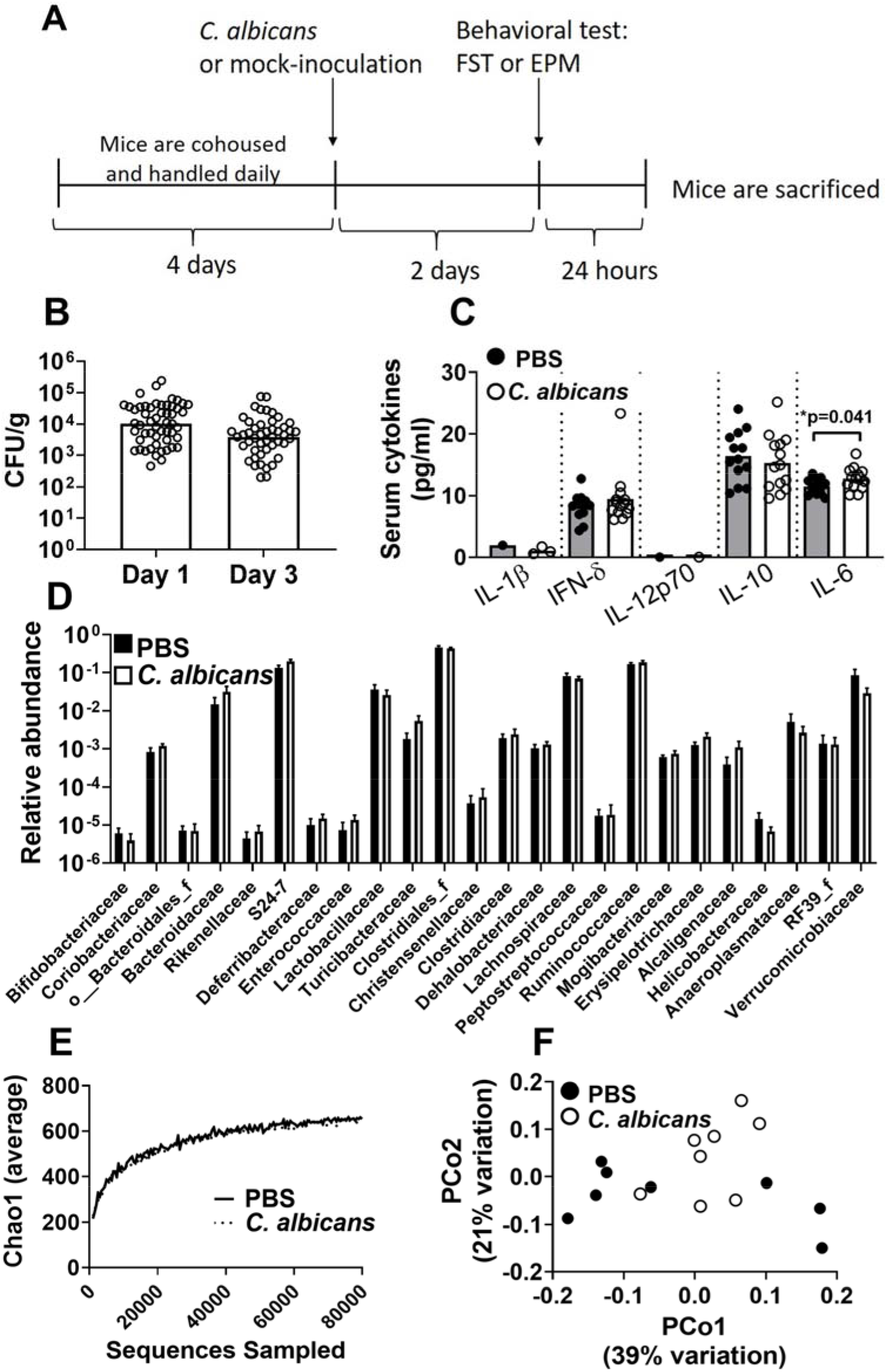
Gastrointestinal colonization with *Candida albicans* after a single inoculation without disruption of bacterial gut microbiota or invasive disease. A) Timeline for acute colonization model. 5-week-old female mice were cohoused for four days and handled daily. Mice were then moved into smaller groups in standard cages and orally inoculated with 5×10^7^CFU of *C. albicans* or given buffer. On day one post-inoculation, fecal pellets were sterilely collected from mice and plated on YPD-SA to measure colonization levels. On day two post-inoculation, all mice underwent a behavioral test. On day three post-inoculation mice were anesthetized with isoflurane and sacrificed by decapitation. Some mice were subjected to restraint stress prior to sacrifice. B) *C. albicans* CFU/g of fecal pellets was measured on day 1 post-inoculation and CFU/g of cecum contents was measured after sacrifice on day three post-inoculation. No culturable fungi were measured on either day from mock-colonized mice. Day 1: N=56, Day 3: N=50. Figure includes data from 6 cohorts. Bars indicate geometric mean. C) A multiplex ELISA was used to measure inflammatory cytokines in the serum of mice. N.D, not detectable. Student's t-test, p<0.05. Mock-colonized N=13, *C. albicans*-colonized N=14. Figure includes data from 3 cohorts. D-F) Microbiota analysis of the cecum contents of mice was performed using standard 16s rRNA DNA sequencing and QIIME analysis pipeline. Mock-colonized N=8 and *C. albicans-colonized* N=8. Figure includes data from 2 cohorts. D) The relative abundance of all bacterial families detected with a median relative abundance greater than 0 is shown. Bar shows geometric mean with SEM E) The average alpha diversity metric Chao1 was calculated to determine the diversity of the microbiota at different levels of sampling. The average Chao1 score of the experimental groups is shown. F) Beta diversity was calculated using weighted UniFrac scores and principal coordinates analysis was performed. For C-D, symbols indicate individual mice and bars indicate the average. Mock-colonized mice are represented as solid dots and bars; *C. albicans-colonized* mice are represented as open circles and bars.

Previous work demonstrated that colonization with *C. albicans* in the absence of antibiotic treatment had a small effect on the composition of the bacterial microbiota (Erb Downward et al., 2013). To determine whether the bacterial microbiota was significantly changed upon *C. albicans* colonization, we analyzed the bacterial microbiota of the cecum of mice sacrificed three days post-inoculation using 16s rRNA DNA sequencing and the QIIME analysis pipeline (Caporaso et al., 2010). The composition and diversity of bacterial taxa were not significantly altered by the introduction of *C. albicans* (Fig. 1D-F), consistent with previous results (Erb Downward et al., 2013). Mann-Whitney U test and linear discriminant analysis effect size ((LEFSE) analysis (Segata et al., 2011) showed that no taxa were significantly different in relative abundance between the microbiota of mock-colonized and *C. albicans-* colonized mice (Fig. 1D). The overall diversity of the microbiota was also not significantly different (Fig. 1E). We analyzed beta diversity and summarized the results using principal coordinates analysis (Fig. 1F). PERMANOVA analysis of the complete principal coordinates analysis found that there was not significant separation of populations based on *C. albicans* colonization (pseudo-F statistic=1.7, p=0.12, 999 permutations). To summarize, mice inoculated with a single dose of *C. albicans* were stably colonized with neither disruption of the bacterial microbiota nor invasive disease.

### 2.2 *C. albicans* colonization increases anxiety-like behavior in the EPM

To assess the effect of GI *C. albicans* colonization on host emotional behavior, standard tests for anxiety-like and stress-coping behavior were used. Two days post-inoculation (Fig. 1A), all mice were subjected to a single behavioral test, either the elevated plus maze (EPM) test for anxiety-like behavior (Walf and Frye, 2007) or the forced swim test (FST) for stress-coping behavior (Hascoët and Bourin, 2009). The EPM consists of two open, aversive arms and two closed arms. Mice were allowed to explore the EPM for a 5-minute trial which was scored by a blinded observer for time spent and entries into the closed and open arms. Anxiety-like behavior is detected as avoidance of the open arms, expressed as percentage arm time (time spent in specified arms divided by the total time spent all arms).

*C. albicans* colonization significantly increased anxiety-like behavior in the EPM. *C. albicans-* colonized mice spent significantly less time in and made fewer entries into the open arms of the EPM compared to the mock-colonized mice (Fig. 2A-B, Mann-Whitney test, p<0.05). There was a significant increase in the percentage of time spent in the closed arms in the *C. albicans*-colonized mice but no difference in the number of entries into the closed arms (Fig. 2C-D). There was no effect of *C. albicans* colonization on the duration of time spent in the center or the total number of entries into all arms (Fig. 2E-F), indicating that *C. albicans* colonization did not alter overall activity but specifically increased anxiety-like behavior in this test.

**Figure 2:**
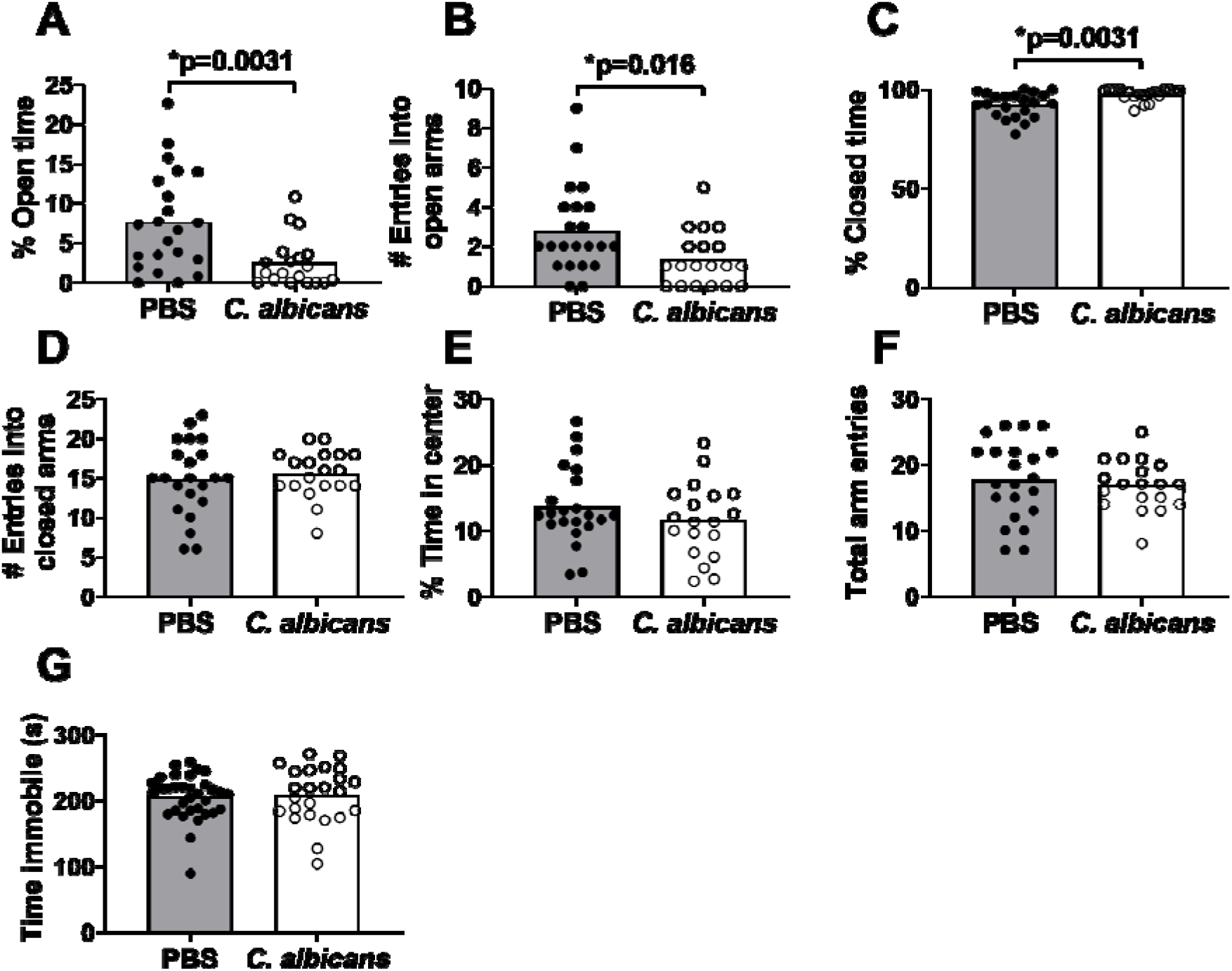
*C. albicans* colonization increases anxiety-like behavior. On day two post-inoculation mice underwent either the Elevated Plus Maze (EPM) or the Forced Swim Test (FST). Trials in the behavioral tests were video-recorded and scored by a blinded observer after the fact. For the EPM data (A-F) that follows, mock-colonized N=22 and *C. albicans-colonized* N=19 and statistical analysis performed was Mann-Whitney U-test (A-C, G) or Student's t-test (D-F), p<0.05. Figures include data from two cohorts. A) Percentage of time spent in the open arms of the EPM. B) Total number of entries into the open arms of the EPM. C) Percentage of time spent in the closed arms of the EPM. D) Total entries into the closed arms of the EPM. E) Percentage of time spent in the neutral central square of the EPM (neither open nor closed arms). F) Total entries during the trial (sum of the open and closed arm entries), a metric for locomotor activity. G) Total time spent immobile during the six-minute FST trial. Immobility was defined as no movements beyond those required to stay afloat. For FST data: mock-colonized N=33, *C. albicans-colonized* N=24. Figure includes data from three cohorts. Throughout, symbols represent individual mice and bars indicate average. Mock-colonized mice are represented as solid dots and bars; *C. albicans-colonized* mice are represented as open circles and bars.

The forced swim test (FST) for stress-coping behavior was used to examine the effect of *C. albicans* colonization on a second aspect of emotional behavior, again two days post-inoculation. Mice were placed in a cylinder containing 22°C water for a six-minute trial which was video-recorded and later scored by a blinded observer for time spent immobile. A passive-coping response in this assay was defined as an increase in the amount of time spent floating immobile rather than the active coping response defined as swimming or actively struggling. There was no difference in the time spent immobile between the *C. albicans*-colonized mice and the mock-colonized mice (Fig. 2G), indicating that *C. albicans* colonization did not affect stress-coping behavior in this test.

### 2.3 *C. albicans* colonization alters production of the stress hormone CORT

The underlying biology of anxiety-like behavior is multifaceted (Nutt et al., 2002). One neuroendocrine pathway that has been shown to be dysregulated in human patients with anxiety is the hypothalamus-pituitary-adrenal (HPA) axis (Kallen et al., 2008). To determine whether *C. albicans* colonization affected the HPA axis, we sacrificed mice three days post-inoculation and measured circulating serum CORT under basal conditions and in response to acute psychological stress.

*C. albicans*-colonized and mock-colonized mice were sacrificed unstressed. *C. albicans-* colonized mice had significantly higher unstressed basal CORT (Fig. 3A Welch's t-test, p=0.0012). When the effect of circadian rhythm on serum CORT was examined, basal CORT in both groups was shown to increase over time throughout the afternoon, as expected (Chung et al., 2017) (Fig. 3B). Basal CORT in *C. albicans-colonized* mice, however, began to rise sooner than in mock-colonized mice and thus was significantly higher in *C. albicans-colonized* mice sacrificed two-to-five hours prior to lights out, consistent with a circadian advance of CORT production (Fig. 3B, ANOVA followed by Sidak's multiple comparison test, p<0.05). These results indicate that *C. albicans* colonization dysregulates circadian control of the HPA axis. Further studies of basal CORT were conducted only with mice sacrificed between 14:00-17:00.

**Figure 3:**
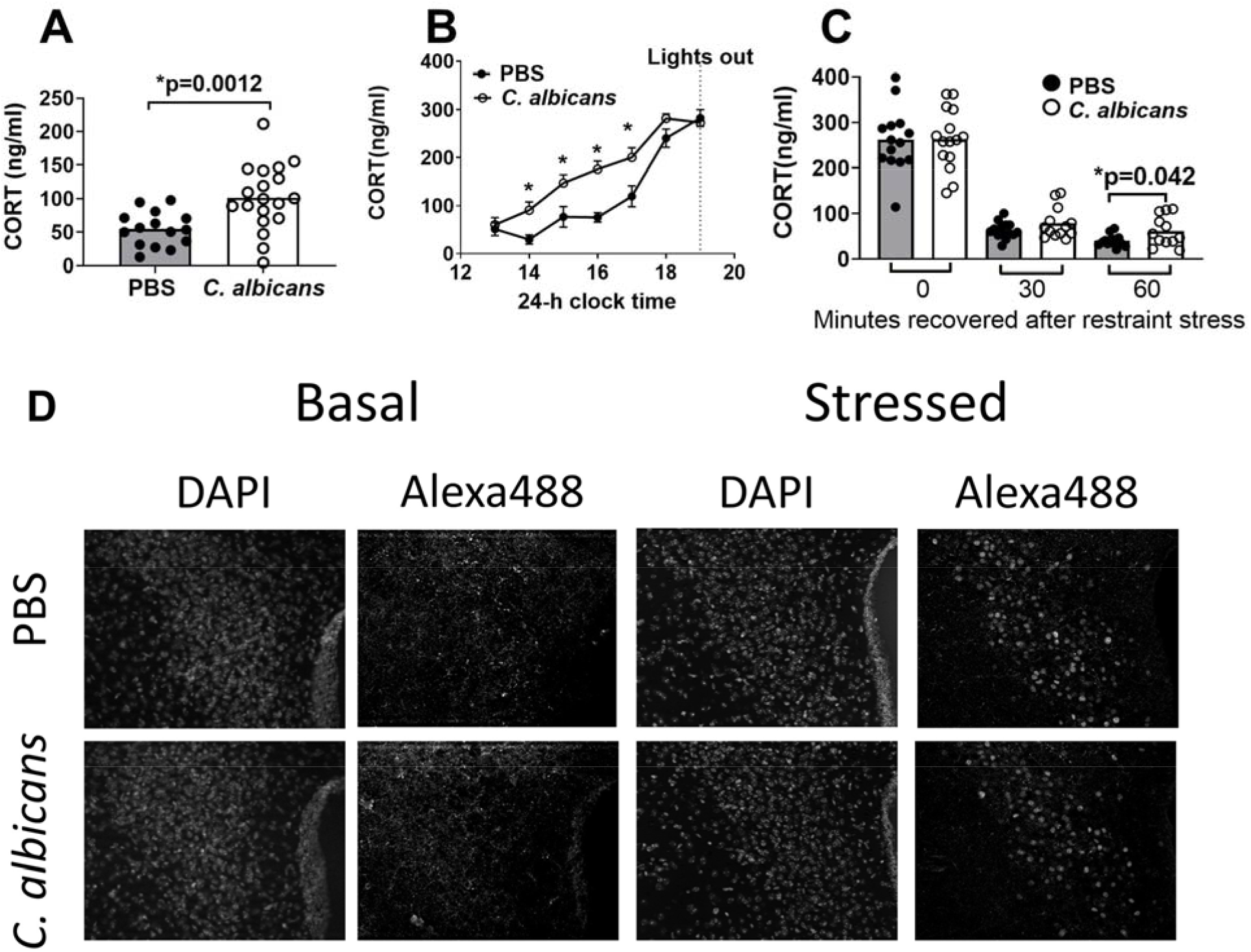
*C. albicans* colonization increases basal CORT and alters feedback inhibition after acute stress. A) Corticosterone (CORT) in trunk blood of mice sacrificed without stress was measured by ELISA. Mock-colonized N=15, *C. albicans*-colonized N=19. Figure includes data from four cohorts. Data was analyzed using Welch's t-test, p<0.05 B) CORT in trunk blood of mice sacrificed at different times of day. In addition to the data summarized in A (mice sacrificed from 13:00-17:00), B includes mice sacrificed without stress from 17:00-20:00. Data was binned in one-hour increments. Mock-colonized: 13:00 N=5, 14:00 N=9, 15:00 N=11, 16:00 N=6, 17:00 N=9, 18:00 N=8, 19:00 N=8. *C. albicans-colonized:* 13:00 N=6, 14:00 N=13, 15:00 N=10, 16:00 N=11, 17:00 N=10, 18:00 N=6, 19:00 N=14. C) Mice were subjected to 30 minutes of restraint stress and then sacrificed immediately (0m recovered) or after 30 minutes or 60 minutes of recovery as shown on the x-axis. Mock-colonized: 0m N= 14, 30m N= 15, 60m N= 13. *C. albicans*-colonized: 0m N= 15, 30m N=13, 60m N= 12. D) Immunohistochemistry was used to visualize cFos protein with Alexa488 in the paraventricular nucleus (PVN) of mice using the 20X objective. DAPI was used to visualize nuclei and Alexa488 to visualize cFOS. The PVN of mice sacrificed unstressed (left) do not exhibit cFOS staining, while PVN of mice sacrificed after 30 minutes of stress and 30 minutes of recovery (right) are positive for cFOS. Representative images are shown. For figures A and C, symbols indicate individual mice and bars indicate the average. For figure B, symbols indicate average and error bars indicate standard deviation. Throughout, mock-colonized mice are solid dots and bars and *C. albicans-colonized* mice are open circles and bars.

To determine whether the changes to basal CORT regulation affected the stress-responsive function of the HPA axis, mice were subjected to 30 minutes of restraint stress and then sacrificed immediately or after 30 minutes or 60 minutes of recovery. *C. albicans*-colonized and mock-colonized mice had comparable peak CORT immediately after stress (Fig. 3C) but *C. albicans-colonized* mice failed to recover from stress to the same extent as the uncolonized mice after 60 minutes of recovery time (Fig. 3C, Student's t-test, p=0.042). These results indicated that *C. albicans* colonization advanced circadian basal CORT production and impaired negative feedback of the HPA axis after stress and recovery.

Activation of the HPA axis begins in the brain, through excitation of the CRH-producing neurons of the paraventricular nucleus (PVN) of the hypothalamus. We used immunofluorescence to quantify expression of cFos protein in the PVN as a marker for neuronal activation. Nuclear cFos protein was not detected in the PVN of unstressed mock-colonized or *C. albicans*-colonized mice (Fig. 3D, left). In mice subjected to 30 minutes of stress and 30 minutes of recovery, nuclear cFos protein was comparably detected in both mock-colonized and *C. albicans*-colonized mice (Fig. 3D, right). The PVN remains stress-responsive in the *C. albicans-colonized* mice, and was not sufficiently stimulated at baseline to be detected by this method. These results demonstrate that *C. albicans* colonization significantly impacts the gut-brain axis through altered basal and stress-induced HPA axis function and increased anxiety-like behavior.

### 2.4 Neuroendocrine changes observed in *C. albicans*-colonized mice are mediated by the endocannabinoid system

The neuroactive lipid endocannabinoid (eCB) N-arachidonoylethanolamide (anandamide; AEA), through its interactions with the CB1 receptor, is a key regulator of anxiety-like behavior and both stress-induced and basal CORT production (Moreira et al., 2008). The circadian advance of basal CORT, reduced feedback inhibition of stress-induced CORT and increased anxiety-like behavior observed in the *C. albicans*-colonized mice resemble the phenotype of the CB1 knock-out mouse (Barna et al., 2004), suggesting a deficit of eCB-CB1 signaling as a result of *C. albicans* colonization. Therefore, we hypothesized that *C. albicans* colonization was interacting with the gut-brain axis via the endocannabinoid system.

To test this hypothesis, we used the well-characterized drug URB597 to increase AEA levels and therefore amplify AEA-CB1 signaling through inhibition of the AEA degradative enzyme, fatty acid amide hydrolase (FAAH) (Fegley et al., 2005, p. 597). Mice were given an intraperitoneal injection of vehicle or URB597 (1mg/kg) 4 hours prior to sacrifice under unstressed conditions, to determine whether elevating AEA levels was sufficient to alleviate the high basal CORT of the *C. albicans-colonized* mice. *C. albicans*-colonized mice given vehicle injection had significantly higher basal CORT than mock-colonized mice as expected (Fig. 4A, Welch's t-test, p=0.0043). Administration of URB597 was sufficient to decrease basal CORT in the *C. albicans-colonized* mice (Fig. 4A, Welch's t-test, p=0.0062). This result is consistent with an AEA deficit in the *C. albicans*-colonized mice underlying the elevated basal CORT. URB597 did not significantly change basal CORT in the mock-colonized mice (Welch's t-test, p=0.10).

**Figure 4:**
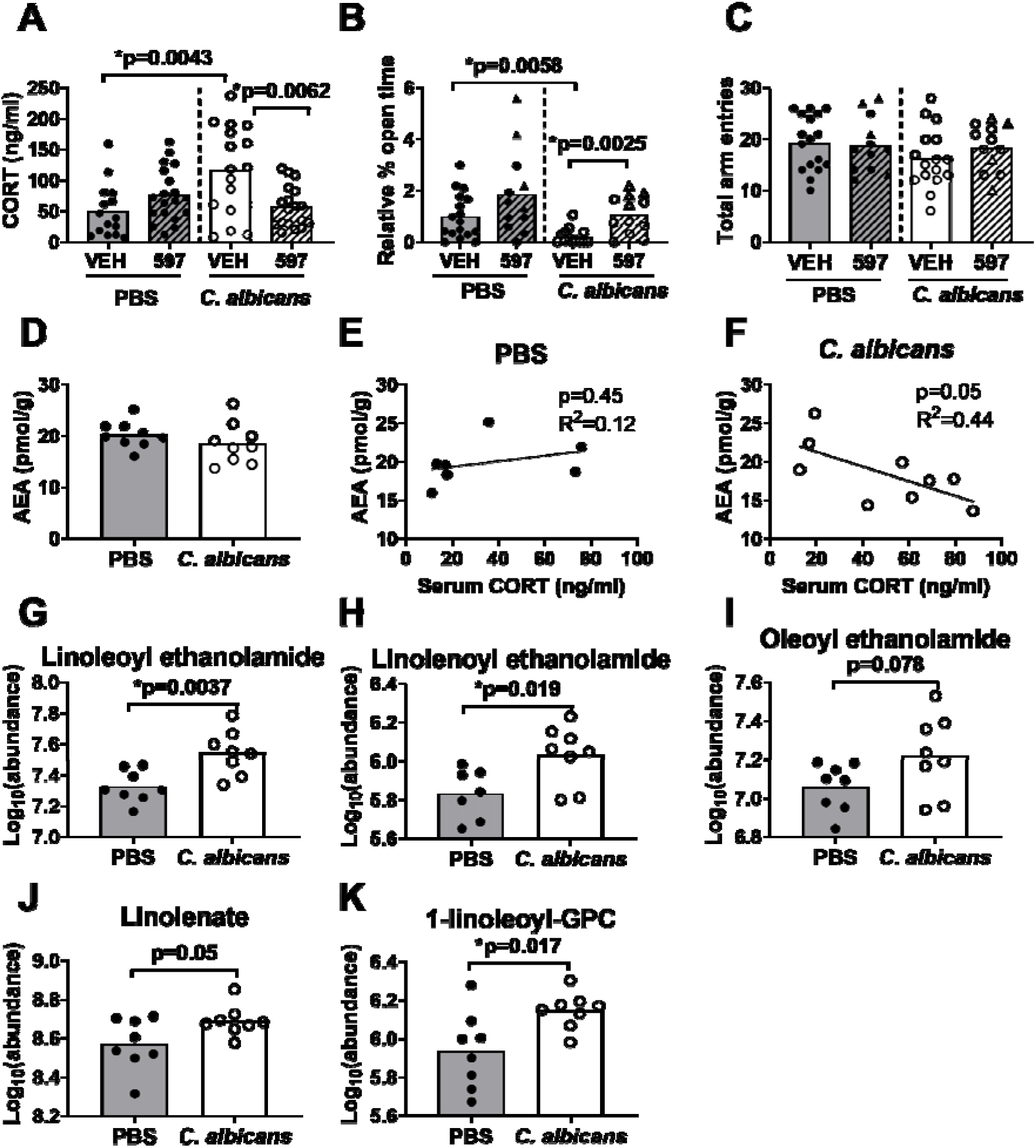
Neuroendocrine phenotypes of the *C. albicans*-colonized mice are mediated through disruption of the endocannabinoid system. A-C) Mice were treated with the fatty acid amide hydrolase (FAAH) inhibitor URB597, a well-characterized drug which increases AEA levels by blocking degradation. Mice were given either URB597 or vehicle control by intraperitoneal injection 4-6 hours prior to sacrifice (A) or testing in the EPM (B-C). A) Mice were treated with 1mg/kg of URB597 or vehicle. ELISA was used to measure CORT in trunk blood of mice sacrificed without stress. Welch's t-test, p<0.05. Mock-colonized: vehicle N=15, URB597 N=19. *C. albicans*-colonized: vehicle N=16, URB597 N=18. Figure includes data from four cohorts. B-C) Mice were treated with 0.1mg/kg (triangles) or 0.15mg/kg (circles) URB597 or vehicle, then tested in the EPM. Mock-colonized: vehicle N=17, 0.1mg/kg URB597 N=4, 0.15mg/kg URB597 N=8. *C. albicans-colonized:* vehicle N=13, 0.1mg/kg URB597 N=4 0.15mg/kg URB597 N=8. B) Percentage time spent in the open arms, normalized to the average percent open time of the mock-colonized vehicle control, is shown. Mann-Whitney U-test, p<0.05. C) Total arms entries, a metric for overall locomotor activity is shown. D) AEA (pmol/g tissue) was measured in forebrain samples by mass spectrometry. E-F) Linear regression of basal serum CORT and forebrain AEA (D) in individual mice are shown. (D-F) Mock-colonized N=7, *C. albicans-colonized* N=9. Untargeted mass spectrometry was used to measure relative abundance of 735 compounds in the cecum contents of mice. N-acylethanolamides (G-I), a free fatty acid (J) and a lysophospholipids (K) containing an 18-carbon chain with varying degrees of unsaturation are shown here. Raw data was log-transformed. Mock-colonized N=8, *C. albicans-colonized*N=8. Student's t-test, p<0.05. Symbols indicate individual mice and bars indicate average. Solid circles and bars are mock-colonized mice and open circles and bars are *C. albicans*-colonized mice.

URB597 was administered 4-6 hours prior to testing in the EPM (0.1-0.15mg/kg) to determine whether increasing AEA-CB 1 signaling was sufficient to alleviate anxiety-like behavior in the *C. albicans-colonized* mice. Due to variability in activity level of mice, data are shown as the percentage of time spent in the open arm normalized to the experimental mean of the mock-colonized vehicle control. As expected, *C. albicans-colonized* mice treated with vehicle had increased anxiety-like behavior compared to the mock-colonized mice (Fig. 4B, Mann-Whitney U-test, p=0.0058). Treatment with URB597 significantly increased open arm time of the *C. albicans*-colonized mice (Fig. 4B, Mann-Whitney U-test, p=0.0025), which indicates that increasing AEA-CB1 signaling was sufficient to normalize anxiety-like behavior in the *C. albicans-colonized* mice. There was no effect of URB597 treatment on overall activity as measured by total arm entries (Fig. 4C). Together, these results demonstrate that increasing AEA-CB1 signaling with the FAAH inhibitor URB597 reversed the effect of *C. albicans* colonization on basal CORT production and anxiety-like behavior, suggesting that the neuroendocrine phenotypes observed in *C. albicans*-colonized mice result from decreased AEA-CB1 signaling.

### 2.5 Basal CORT and brain AEA levels are negatively correlated in *C. albicans*-colonized mice

To test the hypothesis that AEA levels were reduced in colonized mice, AEA levels in the forebrain were measured. Mice were sacrificed on day 3 post-inoculation and forebrain samples analyzed by mass spectrometry. We observed only a trend towards decreased AEA in the forebrains of *C. albicans-colonized* mice compared to mock-colonized mice (Fig. 4D, Student’s t-test, p=0.30). However, a negative correlation between basal serum CORT and AEA in the *C. albicans*-colonized mice (Fig. 4F, R^2^=0.44, F-test, p=0.05) but not in the mock-colonized mice (Fig. 4E, R^2^=0.119, F-test, p=0.45) was observed. Thus mice that responded to *C. albicans* colonization with increased basal CORT did indeed have lower levels of AEA in their forebrain. Previous results have demonstrated a similar negative correlation between basal CORT and AEA levels in the prefrontal cortex of mice (Hill et al., 2010). Taken together, these results indicate that *C. albicans* colonization had variable effects on bulk AEA levels in the mouse forebrain and reduced forebrain AEA was observed in colonized mice with greater dysregulation of the HPA axis.

A second canonical eCB, 2-arachidonoylglycerol (2-AG), was also measured in the forebrain of mice. There was no difference between mock-colonized and *C. albicans*-colonized mice in bulk levels or degree of correlation between 2-AG and serum CORT (Fig. S4).

### 2.6 *C. albicans* colonization alters the gut endocannabinoidome

We performed untargeted metabolomic analysis of the cecum contents of mice to determine whether *C. albicans* colonization altered the metabolite composition of the GI tract and thereby altered eCB metabolism. Of the 735 compounds measured, 21 were significantly differentially abundant in mock-colonized versus *C. albicans-colonized* mice (Table S2, Student's t-test, p<0.05, uncorrected). This result is below the false discovery rate and therefore pathway enrichment analysis was performed to investigate the relevance of the 21 hits. Of the 5 pathways that contained more than one significant compound, two were significantly enriched for *C. albicans-dependent* changes: sterols and eCBs (Fisher's exact test, Holm's correction for multiple comparisons, p<0.0001 and p=0.031 respectively). The enrichment in eCB family compounds indicated that *C. albicans* colonization altered eCB metabolism in the gut, consistent with the eCB changes observed in the brain.

Two such eCB compounds, N-acylethanolamides (NAEs), linoleoyl and linolenoyl ethanolamide, were significantly increased in the *C. albicans-colonized* mice (Fig. 4G-H, Student's t-test, p=0.0037 and p=0.019). An additional NAE, oleoyl ethanolamide, was increased in the *C. albicans-colonized* mice but not significantly (Fig. 4I, Student's t-test, p=0.078). These compounds are structurally similar to anandamide (AEA) but have different acyl groups. AEA itself is present at low levels in the GI tract and was not detected in any of the samples. The increase in 18-C NAEs implies that *C. albicans* colonization altered NAE metabolism. Feeding studies have shown that dietary enrichment for a specific fatty acid (FA) results in overproduction of the NAE derived from the enriched FA and limits production of alternate NAEs (Sihag and Jones, 2019). Membrane phospholipid acyl chains can also act as substrates for the production of NAEs. We measured a trend towards an increase in the 18-C FA linolenate (Fig. 4J, Student's t-test, p=0.05) as well as a significant increase in the lysophospholipid 1-linoleoyl-glycerophosphocholine (Fig. 4K, Student's t-test, p=0.017) in the GI tract of *C. albicans-colonized* mice. Both of these compounds could be converted into their respective 18-C NAEs and increase production of 18C NAEs linoleoyl and linolenoyl ethanolamide while limiting production of AEA. Thus *C. albicans-* induced changes in precursor compound abundance in the GI tract could contribute to the alterations in AEA levels observed in the *C. albicans*-colonized mice.

### 2.7 Altered hepatic lipid metabolism in *C. albicans*-colonized mice reflects gut metabolite changes

In addition to the specific enrichment of two lipid subpathways, sterols and eCBs, and the trend towards increased linolenate described above, we observed an overall increase in lipid compounds containing long-chain polyunsaturated fatty acids (PUFAs) in the *C. albicans-colonized* mice (Fig. S3). Lipids are absorbed from the GI tract into the blood and pass through the liver, a central organ for lipid metabolism which is highly regulated by dietary lipid levels (Xie et al., 2010). We measured liver gene expression to determine whether the changes in lipid levels we quantified in the cecum were sufficient to induce a physiological change in the host.

Mice were sacrificed after three days of *C. albicans* colonization and qRT-PCR was used to measure the expression of lipid-responsive genes in the liver. Decreased expression of the mRNA (*Scd1*) encoding the enzyme stearoyl-coA desaturase (SCD1) was detected in the *C. albicans-colonized* mice compared to the mock-colonized mice (Fig. 5A, Mann-Whitney U-test, p=0.0086), consistent with a transcriptional response to increased PUFA-containing lipids from the GI tract (Lee et al., 1998). Gene expression of regulators of lipogenesis, *Fads1* (acyl-CoA 8-3 desaturase) and *Fads2* (acyl-CoA 6 desaturase), the transcription factor sterol regulatory element-binding protein 1 (SREBP1-c) encoded by *Srebf1* and fatty acid synthase *(Fasn)* (Oosterveer et al., 2009), were not significantly changed by *C. albicans* colonization (Fig. 5B), implying that the increased lipid abundance measured in the cecum was not a result of host lipogenesis but rather that the increased abundance of PUFAs resulted from *C. albicans* colonization and was sufficient to induce hepatic transcriptional changes.

**Figure 5:**
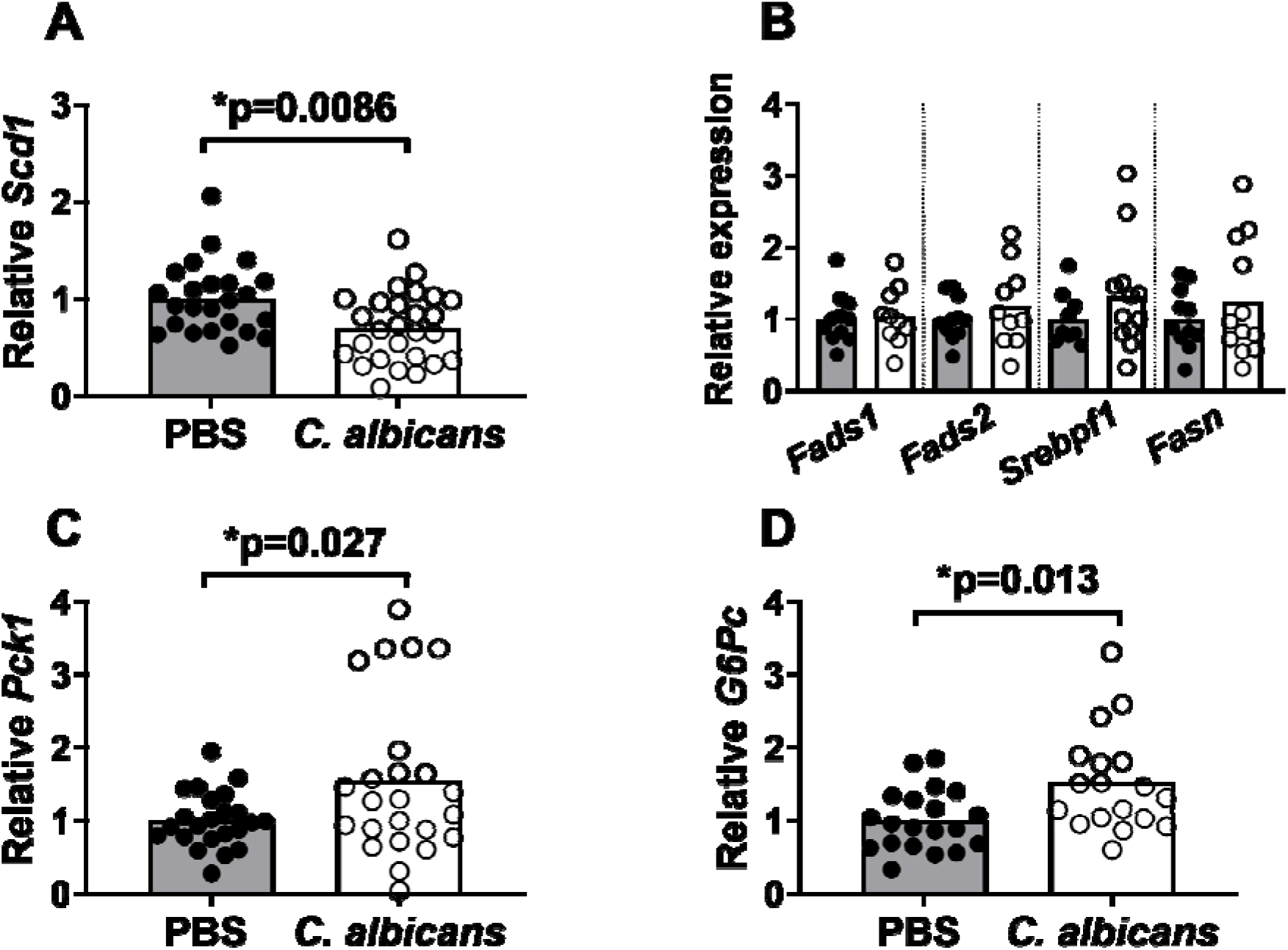
C. albicans colonization affects hepatic lipid-responsive gene expression. RNA was extracted from the liver of mice sacrificed unstressed on day three. cDNA was
synthesized and qRT-PCR was used to quantify liver gene transcription. RNA gene expression is
normalized to GAPDH and expressed relative to the average expression of the mock-colonized mice. A)
Stearoyl-CoA desaturase 1 (*Scd1*) RNA expression in the liver is shown. Mock-colonized N×24, *C. albicans*-colonized N×27. Figure includes data from four cohorts. B) Expression of multiple lipogenic genes is shown: acyl-CoA 8-3 desaturase (*Fads1*), acyl-CoA 6 desaturase (*Fads2*), sterol regulatory element-binding protein 1 (*Srepbf1*) and fatty acid synthase (*Fasn*). Mock-colonized N×11, *C. albicans*-colonized N×10. Figure includes data from two cohorts. C) Expression of the gluconeogenesis enzymes phosphoenolpyruvate carboxykinase (*Pck1*) (mock-colonized N×24, *C. albicans*-colonized N×23) and D) glucose-6-phosphatase (*G6Pc*) is shown (mock-colonized N×20, *C. albicans*-colonized N×18). Figures include data from four cohorts. Symbols represent individual mice and bars indicate average. Solid symbols and bars indicate mock-colonized mice and open symbols and bars indicate *C. albicans*-colonized mice. Mann-Whitney U-test was used for statistical analysis of A and D; Welch’s t-test was used for analysis of B and C, p<0.05.

Also, increased mRNA expression of two enzymes that regulate rate-limiting steps of hepatic gluconeogenesis, phosphoenoylpyruvate carboxykinase (PEPCK-C) encoded by *Pck1* and glucose-6-phosphatase (G6Pase) encoded by *G6pc* was detected in the *C. albicans-colonized* mice compared to the mock-colonized mice (Fig. 5C, Student's t-test, p=0.027, Fig. 5D, Mann-Whitney U-test, p=0.0134). These enzymes can be regulated by both dietary lipid abundance (Massillon et al., 2003) and by basal serum CORT in accordance with the circadian rhythm (Reddy et al., 2007). Thus, the increased expression of gluconeogenic enzymes may reflect both circadian advance of CORT and increased abundance of dietary PUFAs present in *C. albicans-colonized* mice. Regardless, these changes in hepatic gene expression demonstrate that *C. albicans* alters host physiology beyond the GI tract and could have a significant impact on host metabolism.

To summarize, we demonstrated that colonization of mice with the commensal fungus *C. albicans* resulted in significant changes to the gut-brain axis including increased anxiety-like behavior and altered activity of the HPA axis. We identified altered eCB levels in the brain and GI tract of the *C. albicans-colonized* mice and demonstrated that increasing levels of AEA through administration of URB597 was sufficient to reverse the *C. albicans-induced* increase in CORT production and decrease anxiety-like behavior. These results demonstrate that the gut microbiota can alter activity in the brain through modulation of eCB signaling.

## 3.1 Discussion

*C. albicans* is the most common fungal member of the human gut microbiota. We investigated the effect of *C. albicans* colonization on the gut-brain axis and observed specific changes in the endocannabinoidome related to a stress-like behavioral and neuroendocrine phenotype. Previous investigations of the gut-microbiota-brain axis have shown that colonization with commensal bacterial species decreased CORT produced in response to stress and decreased anxiety-like behaviors (Messaoudi et al., 2011; Sudo et al., 2004). In contrast to these bacteria, commensal *C. albicans* increased basal CORT and increased anxiety-like behavior, resembling the neuroendocrine phenotypes observed after infection with the parasite *Trichuris muris* (Bercik et al., 2010) or the mouse gastrointestinal pathogen *Citrobacter rodentium* (Lyte et al., 2006). Researchers demonstrated that infection with these pathogens increased anxiety-like behavior through increased inflammation. *C. albicans* in contrast is a commensal and did not induce significant systemic inflammation. Rather, *C. albicans* colonization changed the gut metabolome and altered eCB signaling to produce the observed increase in basal CORT and anxiety-like behavior. To our knowledge, this is the first communication to report that a commensal fungus, *C. albicans*, affects neuroendocrine host phenotypes and that microbiota-induced changes to eCBs can affect the brain and behavior.

Treatment with the FAAH inhibitor URB597 alleviated both the elevated basal CORT and increased anxiety-like behavior observed in *C. albicans*-colonized mice, indicating a common mechanism of insufficient AEA. Consistent with the results of the URB597 experiments, we measured a trend towards decreased AEA in the forebrain of *C. albicans-colonized* mice, and found a correlation in *C. albicans-colonized* mice such that those with the largest increase in basal CORT also had the lowest AEA, similar to what has been seen in chronic stress models (Hill et al., 2010). This negative correlation supported the model that an AEA deficit was responsible for the neuroendocrine changes observed in *C. albicans-colonized* mice. Previous investigations found that AEA levels specifically within the prefrontal cortex (Hill et al., 2011) and the amygdala (Hill et al., 2010) were responsible for the downregulation of CORT during recovery from stress and for anxiety-like behavior, respectively. It is possible that within those discrete brain regions, colonized mice exhibited a larger decrease in AEA than the decrease we measured in the forebrain as a whole. Untargeted metabolomic analysis of cecum contents further supported the model that *C. albicans* colonization altered eCB metabolism, as two alternate NAEs, linoleoyl and linolenoyl ethanolamide, were increased in abundance in the *C. albicans*-colonized mice as were compounds containing their FA precursors. We suggest that these changes in GI tract eCB levels could reflect systemic changes in lipid availability and NAE production and therefore disruption of normal eCB metabolism. Such disruption could contribute to the altered eCB-CB1 signaling and consequent neuroendocrine changes observed in the *C. albicans*-colonized mice.

Adolescence in humans is a dynamic period of development during which stress and anxiety disorders present a significant health concern (Siegel and Dickstein, 2012); therefore ongoing research into neuroendocrine health using adolescent models is relevant to human health. The current study was performed with mid-adolescent (5-week-old) mice, which were sensitive to HPA axis dysregulation by GI tract colonization with *C. albicans*. Previous work investigating the gut microbiota and the HPA axis demonstrated a window during which mice are sensitive to microbiota manipulation, as Sudo *et al* (2004) found that hyperreactivity of the HPA axis in germ-free mice could be reversed through microbiota transplant in adolescent (6-week-old) mice but not in adult (8-week-old or 14-week-old) mice. Lee *et al* (2013) showed that AEA levels in the brain increased significantly between adolescence and adulthood, and thus the adolescent brain may be especially sensitive to the degree of eCB disruption observed in the *C. albicans-colonized* mice. Previous researchers have also demonstrated significant developmental differences in HPA axis regulation and response in both rodents (Romeo et al., 2014) and humans (Gunnar et al., 2009; Netherton et al., 2004). Women are significantly more likely to experience anxiety disorders than men (McLean et al., 2011), and therefore this study using a female mouse model provides valuable insight into dysregulation of the HPA axis and anxiety-like behavior in a female population.

More broadly, these results highlight the sensitivity of the HPA axis and related anxiety-like behavior to changes in AEA levels, and provide additional evidence that the eCB system is a viable drug target for anxiety disorders. Indeed, a recent study in healthy human volunteers demonstrated that treatment with a different FAAH inhibitor, PF-04457845, raised AEA levels and protected against the development of stress-induced anxiety (Mayo et al., 2019), demonstrating that FAAH inhibition and increased AEA can reverse activation of stress responses and modulate emotional behavior in humans as it does in mouse models. Altogether this work demonstrates the ability of the gut metabolome to have a significant impact on the brain and behavior, and illustrates the ability of the common gut commensal fungus *C. albicans* to affect the host beyond the GI tract.

## 3.2 Conclusions

We have shown that GI colonization with the most prevalent human commensal fungus, *C. albicans*, can alter systemic host health through the gut-brain axis. Although *C. albicans* colonization was limited to the GI tract, mice so colonized exhibited global endocannabinoidome changes that resulted in dysregulation of basal CORT production and increased anxiety-like behavior. We suggest that *C. albicans* is able to induce these changes through manipulation of lipid availability in the GI tract, in agreement with previous studies that have shown the importance of GI lipid pools for eCB production throughout the host (Sihag and Jones, 2019). This study illustrates a novel mechanism by which a member of the microbiota can impact host health, by modulating the GI tract metabolome to change eCB levels throughout the host and alter the gut-brain axis.

## 4. Materials and Methods

Detailed methods are presented in the Supplemental Information

### 4.1 Animals

Up to 23 five-week-old female C57BL/6 mice (Jackson Laboratory) were cohoused in a large cage (24”×17”) and given sterile food, water and bedding. After four days of acclimation, mice were transferred to standard cages with 3-4 mice inoculated with *Candida albicans* or 3-4 mice given a mock-inoculum in each cage. Mice were put through a behavioral test on day two and sacrificed on day three post-inoculation without stress (Sarkar et al., 2011).

All experiments were done in compliance with NIH Guide for the Care and Use of Laboratory Animals and Tufts University IACUC guidelines.

### 4.2 Strains and growth conditions

*C. albicans* strain CKY101 (Brown et al., 1999) was used for all experiments. For preparation of mouse inoculum, cells were grown at 37°C in standard yeast media for 24 hours. 25μl (5×10^7^ cells) of cells in PBS with 2% sucrose was fed to mice for gastrointestinal colonization. 25μl of 2% sucrose in PBS was fed to mice for mock-inoculation.

### 4.3 Drug treatment

URB597 (Sigma Aldrich) and URB937 (Cayman Chemical) were dissolved in 18:1:1 normal saline:PEG400:Tween80 (Sigma Aldrich) and administered to mice via intraperitoneal injection at a dosage of 0.1-1mg/kg bodyweight four hours prior to behavioral testing or sacrifice.

### 4.4 Restraint stress

Mice were placed in a 50ml conical tube with two airholes enclosed with a rubber stopper for 30 minutes. After this restraint, mice were placed back in their home cage and anesthetized and sacrificed or allowed to recover in the home cage for 30 or 60 minutes and then anesthetized and sacrificed.

### 4.5 Elevated Plus Maze

Behavioral testing in the Elevated Plus Maze (EPM) was performed as described in Walf *et al* (2007). After a 5m trial in the EPM, the mouse was placed in a fresh sterile cage. All trials were recorded with a video-camera from above and scored by a blinded observer.

### 4.6 Forced Swim Test

Behavioral testing in the forced swim test was performed as described in Can *et al* (2012). A mouse was removed from its home cage and placed in the beaker of water for a six-minute trial which was recorded by video-camera from the side and scored after the fact by a blinded observer. After testing, mice were placed in a fresh sterile cage.

### 4.7 Bacterial microbiota analysis

The cecum, including contents, was dissected from mice after sacrifice on day three and was immediately frozen on dry ice. Microbial DNA was extracted using the QIAamp DNA Stool Mini Kit (Qiagen) following the manufacturer's protocol. Libraries were prepared from each sample and sequenced as described (Caporaso et al., 2012). The resulting fastq files were used as input for downstream analysis using QIIME (1.8.0)(Caporaso et al., 2010). The resultant OTU tables contained the relative abundance of bacterial taxa in each sample. Analysis of alpha and beta diversity were performed using standard QIIME scripts.

### 4.8 Measurement of hormones and cytokines in serum

Trunk blood was collected into serum separator blood collection tubes (BD) after sacrifice by decapitation. Tubes were then spun to separate serum. Serum was divided into aliquots and frozen at −80°C. Serum corticosterone was measured using a Corticosterone ELISA Kit (Enzo Life Sciences) following the manufacturer's small volume protocol. Manufacturer reported sensitivity down to 27pg/ml of CORT. Authors observed an average intra-assay coefficient of variance of 5.7% and an average interassay coefficient of variance of 8.9%. Serum cytokines were measured using a multiplex ELISA (Quanterix) following manufacturer's protocol. Manufacturer reported sensitivity for each cytokine individually: mIFN-γ 7.1pg/ml, mIL-1β 1.1pg/ml, mIL-6 6.4pg/ml, mIL-10 3.0pg/ml, mIL-12p70 0.37pg/ml. Due to volume of mouse serum required for assay, authors did not have sufficient sample to calculate inter-assay coefficient of variance.

### 4.9 Immunohistochemistry for cFOS in hypothalamus slices

Brains were dissected and fixed in 4% (w/v) paraformaldehyde in PBS for 24h at 4^°^C. They were then cryopreserved in sucrose gradient, rapidly frozen in isopentane chilled on dry ice, and stored at −80°C. Free-floating sections were prepared using a cryostat. Sections of interest were incubated with 1:5000 dilution of rabbit anti-mouse cFos antibody (Sigma Aldrich F7799) for 72h, then stained using anti-rabbit IgG (VectaStain Elite ABC Kit) and streptadvidin-Alexa488 (Molecular Probes). Sections were imaged using a Zeiss microscope with Apotome attachment.

### 4.10 Extraction and measurement of AEA in forebrain

Lipid extraction from forebrain samples was performed as described previously(Morena et al., 2015). AEA was measured using mass spectrometry as described previously (Qi et al., 2015).

### 4.11 Untargeted metabolomic analysis of cecum contents

The cecum was dissected after sacrifice on day three and contents were squeezed into a tube and immediately frozen in dry ice/ethanol bath and stored at −80°C. Extraction of metabolites and untargeted mass spectrometry analysis were performed by Metabolon. Statistical analysis of results was performed using Metabolon Metabolync tools.

### 4.12 Real-time quantitative PCR analysis

Tissues were frozen at −80°C in RNALater (Invitrogen). RNA was purified from tissues using QIAzol for lysis and extraction with a Qialyzer, followed by column purification using the Ambion Purelink Mini kit (Invitrogen). cDNA was synthesized using SuperScript III (Invitrogen) with oligo-dT priming and the manufacturer's protocol. qPCR reactions were performed using SYBR Green Master Mix (Applied Biosystems) and a LightCycler 480 II (Roche) instrument.

### 4.13 Statistical analysis

Statistical anlaysis was performed using GraphPad Prism8.4.0. The majority of the data was analyzed using a two-tailed t-test to compare the means of the two experimental groups in question. When the standard deviation of the data was significantly different between groups, Welch's t-test was used; otherwise Student's t-test was used. Throughout α=0.05 was used to assess significance. When multiple comparisons were required, the Bonferroni correction was used to adjust the value of α. Data was checked for normality using the D'agostino-Pearson omnibus test. When data was not normally distributed, the Mann-Whitney test was used to compare experimental groups.

## 5. Acknowledgments

The authors gratefully acknowledge Dr. Michael Romero for providing critical expertise at an early stage of this project and Dr. Anne Kane for technical assistance in preparation of samples for microbiota analysis. This research was supported by NIH NIAID R01 AI118898 (to C.A.K.). JM is supported by the following grants: R01AA026256, R01NS105628, R01NS105628. LM was also supported by NIH training grant T32AI07422.

