## Supplemental Information for "Colonization with the commensal fungus *Candida albicans* perturbs the gut-brain axis through dysregulation of endocannabinoid signaling"

### Detailed materials and methods

#### Animals

Up to 23, 5-week-old female C57BL/6 mice (Jackson Laboratory) were cohoused in a large cage (24”x17”) and given sterile food, water and bedding. Mice were allowed to acclimate to Tufts Animal Facility for three days during which time they were handled daily. If mice were to be given an intraperitoneal injection of drug/vehicle prior to a behavioral test, mice were also given an intraperitoneal injection (150µl) of sterile normal saline (0.9% w/v) daily during this acclimation period.

The day prior to inoculation with *C. albicans* or mock-inoculation, mice were transferred to standard cages with three or four mice per cage. On this day (day -1), fresh fecal pellets were collected, homogenized in PBS and the homogenate plated on YPD-SA agar (YPD agar plus 100µg/ml streptomycin and 50µg/ml ampicillin) and incubated at 37°C for 2 days to confirm that mice were negative for cultivable fungi. The following day, mice were inoculated orally with 25 µl of *C. albicans* ($5x10^7$CFUs/mouse in PBS with 2% sucrose) or mock-inoculated with 2% sucrose in PBS and moved to a fresh sterile cage (day 0). On day one, fresh fecal pellets were collected from all mice and plated on YPD-SA to measure colonization with *C. albicans*. On day two, mice were put through a behavioral test (either Elevated Plus Maze or Forced Swim), and after the test, transferred from cages of four to cages of two (cages of three were not subdivided) in fresh sterile cages. On day three in most experiments, mice were sacrificed in the afternoon two to four hours prior to lights out. In experiments investigating the effect of the circadian rhythm on CORT production, animals were sacrificed at various other timepoints as noted in the text. Animals were rapidly anesthetized with open-drop exposure to isoflurane and then sacrificed via decapitation.

All experiments were done in compliance with Tufts University IACUC guidelines.

**Strains and growth conditions**

*C. albicans* strain CKY101[1] was used for all experiments. For preparation of mouse inoculum, cells were grown at 37°C in YPD (1% yeast extract (BD), 2% peptone (Difco) 2% glucose (Sigma Aldrich)) for 24 hours. Cells were then washed twice with sterile PBS (phosphate-buffered saline) and resuspended in 2% (w/v) sucrose in PBS at a concentration of $2x10^9$ cells/ml. 25µl ($5x10^7$ cells) of this cell suspension was fed to mice for gastrointestinal colonization. 25µl of a 2% sucrose solution in PBS was fed to mice for mock-inoculation.

**Drug treatment**

URB597 (Sigma Aldrich) and URB937 (Cayman Chemical) were dissolved in 18:1:1 normal saline:PEG400:Tween80 (Sigma Aldrich) and administered to mice via intraperitoneal injection at a dosage of 0.1-1mg/kg bodyweight. Mice received a low dose 4-6 hrs prior to behavioral testing on day 2 and then a higher dose 4-6 hours prior to sacrifice on day 3.

**Restraint stress**

Mice were placed in a 50ml conical tube with two airholes enclosed with a rubber stopper for 30 minutes. After this restraint, mice were placed back in their home cage and either immediately anesthetized and sacrificed or allowed to recover in the home cage for 30 or 60 minutes and then anesthetized and sacrificed.

**Elevated Plus Maze**

Behavioral testing in the Elevated Plus Maze (EPM) was performed as described in Walf *et al* 2013[2]. Mice were moved from their housing room to the testing room at least one hour prior to testing. The EPM was sprayed with 70% ethanol and thoroughly dried. A mouse was then removed from its home cage, placed in the center of the EPM facing an open arm and allowed to explore for five minutes. After testing the mouse was placed in a fresh sterile cage. The EPM was sprayed with 70% ethanol prior to each trial. All trials were recorded with a video-camera from above and scored after the fact by a blinded observer for the number of open arm entries and the duration of time spent in the open arms. A mouse was scored as having entered an open arm when all four paws crossed into the open arm

**Forced Swim Test**

Behavioral testing in the forced swim test was performed as described in Can et al 2012 [3]. Mice were moved from their housing room to the testing room at least one hour prior to testing. An 8.3L cylinder was sprayed with disinfectant and wiped dry, then filled with 10cm of 22°-25°C tap water. A mouse was removed from its home cage and placed in the beaker of water for a six-minute trial which was recorded by video-camera from the side and scored after the fact by a blinded observer. Trials were monitored to ensure mice remained floating throughout. After testing, mice were placed in a fresh sterile cage. FST videos were manually scored for the number of seconds spent floating immobile.

**Bacterial microbiota analysis**

The cecum, including contents, was dissected from mice after sacrifice on day three and was immediately frozen on dry ice. Microbial DNA was extracted using the QIAamp DNA Stool Mini Kit (Qiagen) following the manufacturer's protocol. Briefly, cecum samples were lysed by beadbeating (samples combined with 500mg 0.1mm diameter zirconia/silica beads) in Qiagen lysis buffer ASL, and the lysate was treated with InhibitEX tablets followed by enzymatic digestion with proteinase K (>500mAU/ml) and RNaseA (1mg/ml) and column DNA purification. Libraries were prepared from each sample and sequenced as described[4]. Briefly, PCR amplification of the V4 region of the 16S rRNA gene was performed with primers that included adapters for Illumina sequencing and twelve base barcodes. Two hundred fifty bp paired-end sequencing was performed using an Illumina MiSeq. Base calling was performed using CASAVA 1.8 and the resulting fastq files were used as input for downstream analysis using QIIME (1.8.0)[5]. Briefly, the paired-end reads from the fastq files were joined, barcodes extracted and then demultiplexed. The operational taxonomic units (OTUs) were determined using a closed reference approach by aligning reads to the Greengenes Database (version 13_8) at 99% identity. The Greengenes phylogenetic tree was used to define the phylogenetic relationship between OTUs. The resultant OTU tables contained the relative abundance of bacterial taxa in each sample. These tables were used to calculate overall diversity within each sample. To compare the composition and diversity of samples to each other taking into account phylogenetic relatedness, the OTU tables were used to calculate the weighted UniFrac distance matrix, which was summarized with Principal Coordinate Analysis. Permanova analysis was performed using QIIME.

**Measurement of hormones and cytokines in serum**

Trunk blood was collected into serum separator blood collection tubes (BD) after sacrifice by decapitation and allowed to sit at room temperature for up to three hours. Tubes were then spun at 10,000xg for 10 minutes to separate serum. Serum was divided into aliquots and frozen at -80°C.

Serum corticosterone was measured using a Corticosterone ELISA Kit (Enzo Life Sciences) following the manufacturer's small volume protocol including the serum displacement reagent. Serum cytokines were measured using a multiplex ELISA (Quanterix) following manufacturer's protocol.

**Immunohistochemistry for cFOS in hypothalamus slices**

Brains were rapidly dissected and fixed in 4% (w/v) paraformaldehyde in PBS for 24h at 4°C. They were then cryopreserved by incubation in 10% (w/v) sucrose in PBS for 24h followed by 24h incubation in 30%(w/v) sucrose in PBS. Brains were rapidly frozen in isopentane chilled on dry ice and stored at -80°C. They were sectioned into 40μm sections using a cryostat and gross anatomical landmarks were used to identify sections containing the PVN of the hypothalamus. Sections of interest were incubated with 1:5000 dilution of rabbit anti-mouse cFos antibody (Sigma Aldrich F7799) for 72h, then stained using biotinylated anti-rabbit IgG (VectaStain Elite ABC Kit) and streptadvidin-Alexa488 (Molecular Probes). Sections were then mounted onto glass slides and imaged using a Zeiss microscope with Apotome attachment. Comparable results were observed in a limited study using a Leica SP8 confocal microscope.

**Untargeted metabolomic analysis of cecum contents**

The cecum was dissected after sacrifice on day three and contents were squeezed into a tube and immediately frozen in dry ice/ethanol bath and stored at -80°C. Extraction of metabolites and untargeted metabolomic analysis were performed by Metabolon. As per their report: samples were prepared using the automated MicroLab STAR® system from Hamilton Company. To remove protein, dissociate small molecules bound to protein or trapped in the precipitated protein matrix, and to recover chemically diverse metabolites, proteins were precipitated with methanol under vigorous shaking for 2 min (Glen Mills GenoGrinder 2000) followed by centrifugation. The resulting extract was divided into five fractions: two for analysis by two separate reverse phase (RP)/UPLC-MS/MS methods with positive ion mode electrospray ionization (ESI), one for analysis by RP/UPLC-MS/MS with negative ion mode ESI and one for analysis by HILIC/UPLC-MS/MS with negative ion mode ES (additional fraction saved as back-up). All methods utilized a Waters ACQUITY ultra-performance liquid chromatography (UPLC) and a Thermo Scientific Q-Exactive high resolution/accurate mass spectrometer interfaced with a heated electrospray ionization (HESI-II) source and Orbitrap mass analyzer operated at 35,000 mass resolution. The MS analysis alternated between MS and data-dependent $MS^n$ scans using dynamic exclusion. The scan range varied slighted between methods but covered 70-1000 m/z. Raw data was extracted, peak-identified and QC processed using Metabolon's hardware and software.

Mass spectrometry data was log-transformed and statistical analysis of the results was performed using MetaboAnalyst 4.0[6,7].

**Real-time quantitative PCR analysis**

For measurement of gene expression, mice were sacrificed on day three and tissues were frozen at -80°C in RNALater (Invitrogen). RNA was purified from tissues using QIAzol for lysis and extraction with a Qialyzer, followed by column purification using the Ambion Purelink Mini kit (Invitrogen). cDNA was synthesized using SuperScript III (Invitrogen) with oligo-dT priming and the manufacturer's protocol. qPCR reactions were performed using SYBR Green Master Mix (Applied Biosystems) and a LightCycler 480 II (Roche) instrument. Standard curves were generated and all results normalized to the level of GAPDH expression in each sample. All primers are listed in Table S2. Expression levels were normalized to the experimental mean of the control group.

**Lipid extraction and endocannabinoid analysis of forebrain samples**

Lipid extractions were carried out as previously described[8]. Briefly, frozen brain tissue was weighed and then manually homogenized (with a glass rod) in borosilicate glass culture tubes containing 2ml of acetonitrile with 5 nmol of d8-2-AG, 5 pmol of d8-AEA, 40 pmol d4-PEA, and 40 pmol d4-OEA. All samples were sonicated for 30min in an ice bath and incubated overnight at -20°C to precipitate proteins. The following day samples were centrifuged at 1500xg to remove particulates. The supernatant from each sample was transferred to a new glass tube and evaporated under nitrogen, the tube was then washed once with 350µl acetonitrile (to recapture any lipids adhering to the glass wall) and the acetonitrile was dried under nitrogen gas again. After completely drying, the samples were re-suspended in 200µl of acetonitrile and stored at -80°C until analysis by liquid chromatography mass spectrometry. Analysis in mass spectrometry was performed exactly as previously described[9].

**Supplemental materials and methods references**

**Supplemental figures**

**Figure S1**

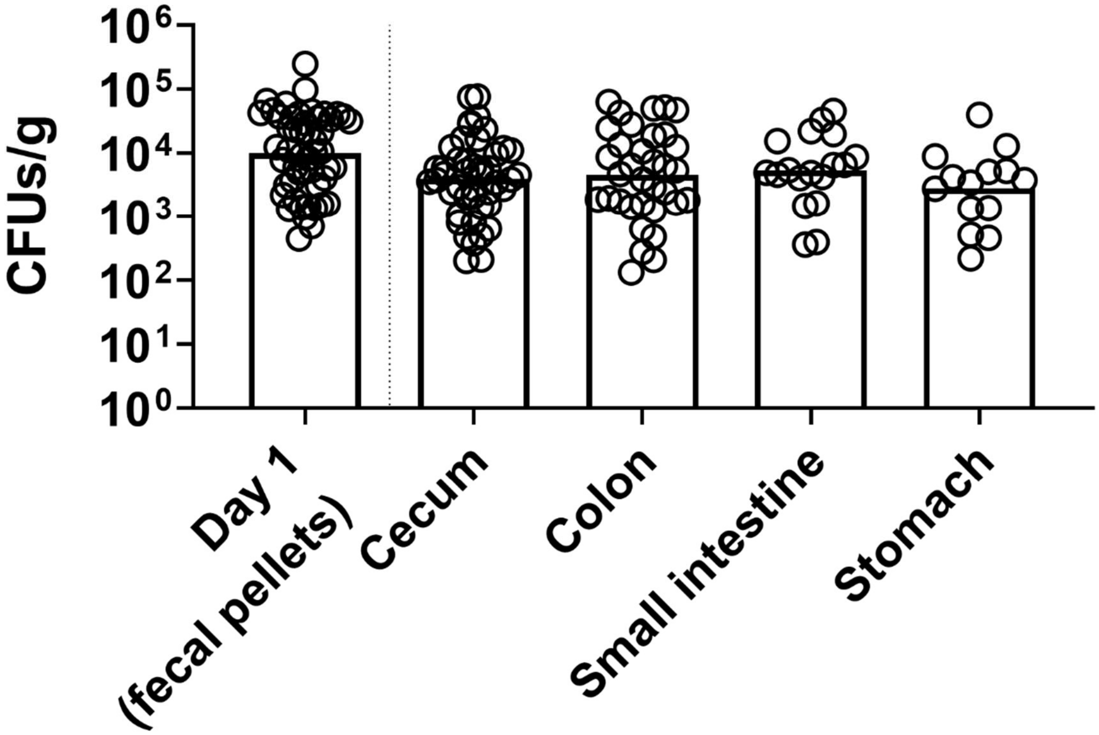


**Figure S1: *C. albicans* CFU/g measured from different sections of the GI tract after sacrifice.** Left: fecal pellets were collected into sterile PBS and homogenate plated on YPD-SA to measure *C. albicans* colonization load. Right: Mice were sacrificed on day three post inoculation and the contents of various organs was collected into sterile PBS and the homogenate plated on YPD-SA to measure the *C. albicans* colonization level of each organ, quantified as colony-forming units (CFU) per gram of material. Each symbol indicates an individual mouse, bars indicate the geometric mean. Figure summarizes results from 2-3 cohorts (varies based on organ).

**Figure S2**

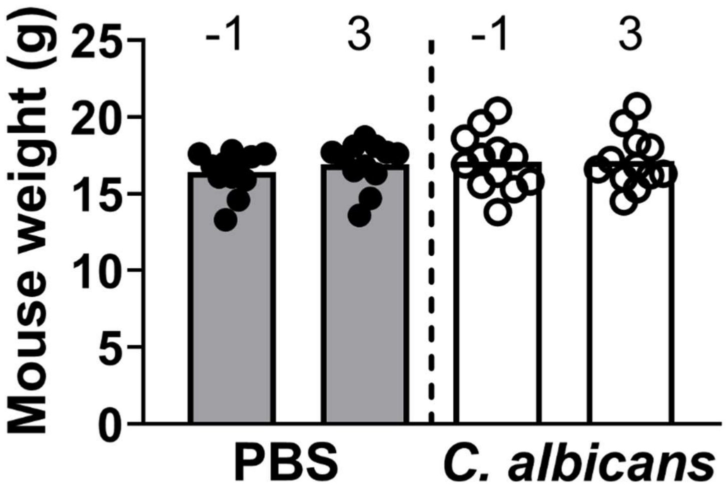


**Figure S2: Mouse weight over time.** Mice were weighed prior to inoculation or mock-inoculation (day -1) and weighed again prior to sacrifice (day 3). No mouse in either group lost >5% of its starting bodyweight. Symbols indicate individual mice and bars indicate the average value of the experimental group.

**Figure S3**

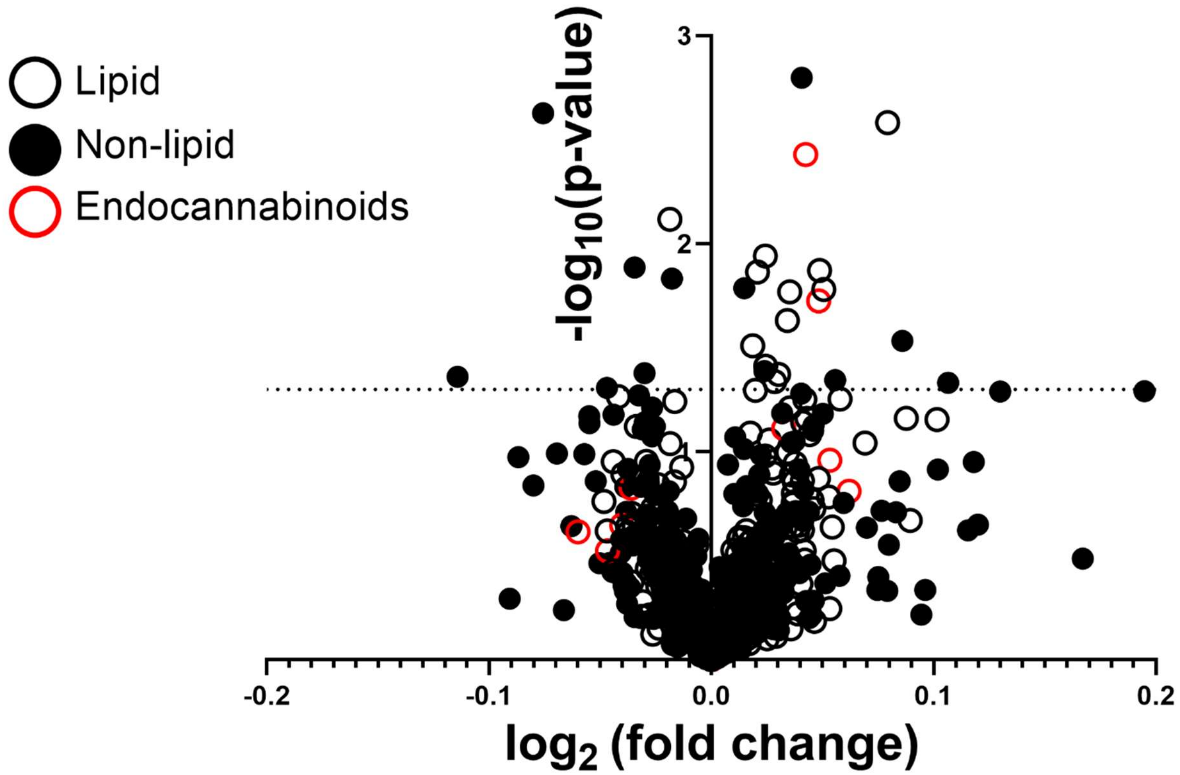


**Figure S3: Untargeted metabolomics analysis of cecum contents.** Cecum contents from 8 uncolonized mice and 8 *C. albicans*-colonized mice were analyzed using mass spectrometry by Metabolon. This figure shows all 735 compounds detected in the study as a volcano plot with the $\log_2$(fold change) along the x-axis and the $-\log_{10}$(p-value) of an unpaired t-test (without additional correction) along the y-axis. Lipid compounds are shown as open circles and endocannabinoids (a subclass of lipids) are shown as red open circles. All other classes of compounds are shown as solid circles. The dotted line at y=1.3 indicates an uncorrected significance cutoff of p=0.05.

**Figure S4**

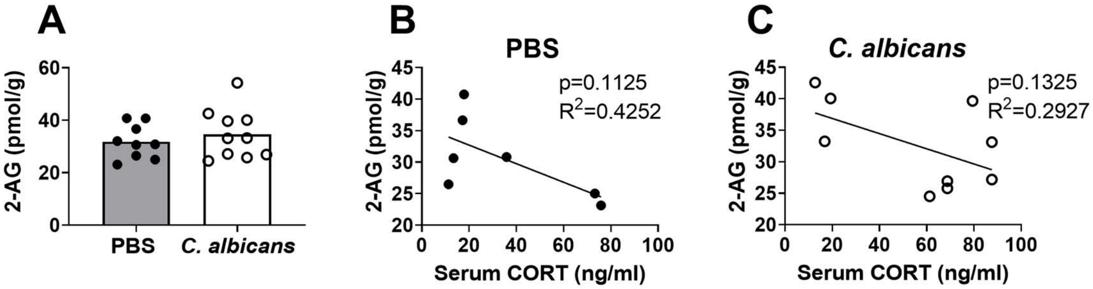


**Figure S4: Forebrain 2-AG levels are not affected by *C. albicans* colonization and do not significantly correlate with serum CORT.** Mice were sacrificed without stress after three days of colonization with *C. albicans* or post-mock-colonization. Lipids were extracted from whole forebrain samples and levels of the endocannabinoids AEA (Fig. 4D-F) and 2-arachidonoylglycerol (2-AG) were measured using mass spectrometry. A) Total bulk level of 2-AG measured in the forebrain of mice. B-C) Correlation between 2-AG and basal serum CORT in either mock-colonized (B) and *C. albicans*-colonized (C) mice. Symbols represent individual mice and bars indicate the average. B and C show a linear regression, the $R^2$ value indicates goodness of fit and the p-value the F statistic. Mock-colonized PBS N=7 (closed symbols) and *C. albicans*-colonized N=9 (open symbols).

**Table S1**

| Gene | Primer sequence | Reference |
|---|---|---|
| *Scd1*, stearoyl-CoA desaturase | F- AGCTCAGTCTCACTCCTTCCCTTA<br>R- CAGCCAGCCTCTTGACTATTCC | Kajikawa et al, 2009[61] |
| *Fads1*, acyl-CoA 8-3 desaturase | F- CCAGCTTTGAACCCACCAA<br>R- CATGAGGCCCATTCGCTCTA | Su et al, 2016[62] |
| *Fads2*, acyl-CoA 6 desaturase | F-TCAAAACCAACCACCTGTTCTTC<br>R-GATGAACCAGGCAAGGCTTTC | Rosenblat et al, 2010[63] |
| *Srebpf1*, sterol regulatory element binding-protein 1 | F- CGGCGCGGAAGCTGT<br>R- TGCAATCCATGGCTCCGT | Kajikawa et al, 2009[61] |
| *Fasn*, fatty acid synthase | F- TACCAGTGCCACAGGAGTCTCA<br>R- TAAACACCTCGTCGATTTCGTTC | Osei-Hyiaman et al, 2005[64] |
| *Pck1*, phosphoenoylpyruvate carboxykinase | F- GTGCTGGAGTGGATGTTCGG<br>R- CTGGCTGATTCTCTGTTTCAGG | Volat et al, 2012[65] |
| *G6Pc*, glucose-6-phosphatase | F- ACTGTGGGCATCAATCTCCTC<br>R- CGGGACAGACAGACGTTCAGC | Volat et al, 2012[65] |
| *Gapdh*, glyceraldehyde-3-phosphate dehydrogenase | F- TGTAGACCATGTAGTTGAGGTCA<br>R- AGGTCGGTGTGAACGGATTTG | Overbergh et al, 1999[66] |

**Table S1: Primer sequences used for qRT-PCR.** This table includes all primer sequences for the qRT-PCR data described in this study. Expression of genes of interest was normalized to GAPDH.

**Table S2**

**Table S2: All compounds detected in untargeted metabolomics study.** This table includes all 735 compounds detected in the 16 mice included in this study. The identity of each compound and the pathway to which that compound was assigned was provided by Metabolon. Data was log-transformed and statistical analysis performed by researcher. Table displays the p-value of an unpaired t-test comparing the abundance of each compound in *C. albicans*-inoculated to mock-inoculated mice (not corrected for multiple comparisons) and the fold change ratio of the abundance of the compounds in the *C. albicans*-inoculated group compared to the mock-inoculated group.

| Biochemical compound | Pathway | P-value (t-test) | Fold Change |
|---|---|---|---|
| asparagine | Amino Acid | 0.227912 | 0.957311935 |
| N-acetylalanine | Amino Acid | 0.436513 | 1.00764636 |
| N-acetylasparagine | Amino Acid | 0.454163 | 0.989420426 |
| alanine | Amino Acid | 0.504343 | 0.997921886 |
| N-acetylaspartate (NAA) | Amino Acid | 0.521837 | 0.987217128 |
| aspartate | Amino Acid | 0.708585 | 0.9982889 |
| creatinine | Amino Acid | 0.673862 | 1.008835958 |
| creatine | Amino Acid | 0.718031 | 1.010023586 |
| guanidinoacetate | Amino Acid | 0.892881 | 1.004393258 |
| N-acetylglutamine | Amino Acid | 0.012994 | 0.976312103 |
| N-methyl-GABA | Amino Acid | 0.067573 | 0.962569379 |
| pyroglutamine* | Amino Acid | 0.154605 | 0.986816264 |
| carboxyethyl-GABA | Amino Acid | 0.206922 | 0.980698356 |
| alpha-ketoglutaramate* | Amino Acid | 0.283307 | 0.986240832 |
| glutamate | Amino Acid | 0.317979 | 0.995291878 |
| N-acetylglutamate | Amino Acid | 0.337511 | 0.989199753 |
| gamma-aminobutyrate (GABA) | Amino Acid | 0.4213 | 0.99367546 |
| beta-citrylglutamate | Amino Acid | 0.696037 | 1.016650394 |
| S-1-pyrroline-5-carboxylate | Amino Acid | 0.74943 | 0.995589615 |
| glutamate, gamma-methyl ester | Amino Acid | 0.799112 | 0.997249791 |
| glutamine | Amino Acid | 0.99129 | 1.000105015 |
| 2-hydroxybutyrate/2-hydroxyisobutyrate | Amino Acid | 0.066417 | 0.969966621 |
| 2-aminobutyrate | Amino Acid | 0.267446 | 0.980338167 |
| 5-oxoproline | Amino Acid | 0.370414 | 0.993908487 |
| cysteinylglycine | Amino Acid | 0.415928 | 1.022855832 |
| cysteinylglycine disulfide* | Amino Acid | 0.904375 | 0.996850581 |
| dimethylglycine | Amino Acid | 0.14531 | 0.946031646 |
| betaine | Amino Acid | 0.307272 | 0.985625727 |
| sarcosine | Amino Acid | 0.35395 | 1.009282473 |

| Biochemical compound | Pathway | P-value (t-test) | Fold Change |
|---|---|---|---|
| N-acetylthreonine | Amino Acid | 0.401278 | 0.98889738 |
| N-acetylglycine | Amino Acid | 0.612132 | 0.99692934 |
| O-acetylhomoserine | Amino Acid | 0.629828 | 1.004411112 |
| N-acetylserine | Amino Acid | 0.652708 | 0.994399391 |
| glycine | Amino Acid | 0.662126 | 0.997763849 |
| threonine | Amino Acid | 0.701316 | 1.001583413 |
| serine | Amino Acid | 0.70976 | 0.998516695 |
| 4-guanidinobutanoate | Amino Acid | 0.536221 | 0.990384241 |
| N-acetyl-1-methylhistidine* | Amino Acid | 0.238649 | 1.083302805 |
| N-acetylhistamine | Amino Acid | 0.290073 | 1.009602499 |
| N-acetylhistidine | Amino Acid | 0.357965 | 0.986614959 |
| imidazole lactate | Amino Acid | 0.380958 | 0.983312604 |
| 4-imidazoleacetate | Amino Acid | 0.406735 | 1.01983583 |
| 1-methyl-4-imidazoleacetate | Amino Acid | 0.44165 | 1.006752445 |
| N-acetyl-3-methylhistidine* | Amino Acid | 0.473422 | 1.022423598 |
| hydantoin-5-propionate | Amino Acid | 0.500761 | 1.012852604 |
| histamine | Amino Acid | 0.523448 | 1.008959742 |
| histidine | Amino Acid | 0.559651 | 0.996054932 |
| formiminoglutamate | Amino Acid | 0.677759 | 1.007184737 |
| imidazole propionate | Amino Acid | 0.752943 | 1.009970384 |
| cis-urocanate | Amino Acid | 0.7896 | 1.003663662 |
| anserine | Amino Acid | 0.867352 | 1.001915632 |
| trans-urocanate | Amino Acid | 0.916343 | 0.998706189 |
| 3-methyl-2-oxobutyrate | Amino Acid | 0.041846 | 0.979357347 |
| ethylmalonate | Amino Acid | 0.049327 | 0.967939882 |
| isobutyrylglycine | Amino Acid | 0.052611 | 1.028385543 |
| 3-methyl-2-oxovalerate | Amino Acid | 0.06121 | 0.981677597 |
| 4-methyl-2-oxopentanoate | Amino Acid | 0.084441 | 0.981697087 |
| 1-carboxyethylisoleucine | Amino Acid | 0.30662 | 0.972129056 |
| N-acetylleucine | Amino Acid | 0.308319 | 0.985831413 |
| N-acetylisoleucine | Amino Acid | 0.330856 | 0.983964362 |
| 2,3-dihydroxy-2-methylbutyrate | Amino Acid | 0.348871 | 1.031394396 |
| 3-methylglutaconate | Amino Acid | 0.403773 | 0.984982212 |
| isovalerylglycine | Amino Acid | 0.417773 | 1.009128779 |
| 1-carboxyethylleucine | Amino Acid | 0.588096 | 1.015387538 |
| 2-hydroxy-3-methylvalerate | Amino Acid | 0.630151 | 0.986255334 |
| leucine | Amino Acid | 0.667661 | 0.998587973 |
| valine | Amino Acid | 0.735115 | 0.998491601 |
| methylsuccinate | Amino Acid | 0.739672 | 0.997032094 |
| isoleucine | Amino Acid | 0.75782 | 0.998938843 |

| Biochemical compound | Pathway | P-value (t-test) | Fold Change |
|---|---|---|---|
| 3-hydroxy-2-ethylpropionate | Amino Acid | 0.761788 | 0.996747774 |
| isovalerate (i5:0) | Amino Acid | 0.763873 | 1.004734077 |
| N-acetylvaline | Amino Acid | 0.797918 | 1.003001894 |
| alpha-hydroxyisovalerate | Amino Acid | 0.804881 | 0.99348774 |
| alpha-hydroxyisocaproate | Amino Acid | 0.826648 | 0.995024493 |
| 3-hydroxyisobutyrate | Amino Acid | 0.974996 | 1.0011298 |
| beta-hydroxyisovalerate | Amino Acid | 0.982036 | 0.9997862 |
| N2,N6-diacetyllysine | Amino Acid | 0.138903 | 0.964581721 |
| N6-acetyllysine | Amino Acid | 0.202994 | 0.980268644 |
| pipecolate | Amino Acid | 0.310266 | 0.992459196 |
| N,N,N-trimethyl-5-aminovalerate | Amino Acid | 0.343916 | 0.965708737 |
| 2-oxoadipate | Amino Acid | 0.376982 | 0.985512179 |
| N2-acetyllysine | Amino Acid | 0.388653 | 0.987421331 |
| 2-aminoadipate | Amino Acid | 0.400079 | 1.004754335 |
| 5-aminovalerate | Amino Acid | 0.498632 | 0.99639285 |
| N-acetyl-cadaverine | Amino Acid | 0.545347 | 0.994906182 |
| N6,N6,N6-trimethyllysine | Amino Acid | 0.614566 | 1.005245989 |
| saccharopine | Amino Acid | 0.677578 | 0.989571636 |
| lysine | Amino Acid | 0.680754 | 0.998387466 |
| cysteine s-sulfate | Amino Acid | 0.079163 | 1.032197556 |
| alpha-ketobutyrate | Amino Acid | 0.106129 | 0.941567255 |
| 4-methylthio-2-oxobutanoate | Amino Acid | 0.134715 | 0.980592946 |
| 3-sulfo-L-alanine | Amino Acid | 0.243185 | 0.979159771 |
| methionine sulfoxide | Amino Acid | 0.262258 | 0.995946282 |
| cysteine sulfinic acid | Amino Acid | 0.297983 | 1.011661029 |
| N-acetyltaurine | Amino Acid | 0.371461 | 0.981812868 |
| cystine | Amino Acid | 0.427654 | 0.990908872 |
| N-acetylcysteine | Amino Acid | 0.449707 | 0.986101538 |
| methionine | Amino Acid | 0.460728 | 0.997020344 |
| cysteine | Amino Acid | 0.611931 | 0.990276703 |
| taurine | Amino Acid | 0.667592 | 1.006323579 |
| N-formylmethionine | Amino Acid | 0.975 | 0.99978312 |
| N-acetylmethionine sulfoxide | Amino Acid | 0.981467 | 0.999511971 |
| N-acetylmethionine | Amino Acid | 0.983541 | 1.000302936 |
| phenylpyruvate | Amino Acid | 0.078151 | 0.978784079 |
| 4-hydroxyphenylacetate | Amino Acid | 0.468224 | 1.02133497 |
| phenylalanine | Amino Acid | 0.750908 | 0.99889735 |
| N-acetylphenylalanine | Amino Acid | 0.753353 | 1.003702537 |
| phenylacetate | Amino Acid | 0.827705 | 1.005513385 |
| phenyllactate (PLA) | Amino Acid | 0.835652 | 1.003178084 |

| Biochemical compound | Pathway | P-value (t-test) | Fold Change |
|---|---|---|---|
| 3-hydroxyphenylacetate | Amino Acid | 0.838655 | 0.99883802 |
| 1-carboxyethylphenylalanine | Amino Acid | 0.987306 | 0.999568654 |
| N-acetyl-isoputreanine* | Amino Acid | 0.19284 | 1.026634614 |
| diacetylspermidine* | Amino Acid | 0.194743 | 0.973773003 |
| N-acetylputrescine | Amino Acid | 0.275031 | 1.018600527 |
| spermidine | Amino Acid | 0.381856 | 0.972673965 |
| (N(1) + N(8))-acetylspermidine | Amino Acid | 0.698211 | 0.994689293 |
| 4-acetamidobutanoate | Amino Acid | 0.792197 | 1.002471976 |
| N1,N12-diacetylspermine | Amino Acid | 0.938411 | 0.996605337 |
| indole-3-carboxylate | Amino Acid | 0.001594 | 1.028581099 |
| kynurenate | Amino Acid | 0.390944 | 1.021120625 |
| 5-hydroxyindoleacetate | Amino Acid | 0.421937 | 0.990174696 |
| indole | Amino Acid | 0.444979 | 0.974429657 |
| N-acetyltryptophan | Amino Acid | 0.549895 | 1.009449183 |
| indolepropionate | Amino Acid | 0.567403 | 0.992161496 |
| serotonin | Amino Acid | 0.591553 | 1.008402126 |
| indolelactate | Amino Acid | 0.682959 | 1.00561252 |
| tryptophan | Amino Acid | 0.730407 | 1.00144837 |
| xanthurenate | Amino Acid | 0.755753 | 1.006919028 |
| indoleacetate | Amino Acid | 0.82096 | 1.002309837 |
| N-formylanthranilic acid | Amino Acid | 0.837529 | 0.999081326 |
| kynurenine | Amino Acid | 0.845403 | 0.995715211 |
| picolinate | Amino Acid | 0.984911 | 0.999878485 |
| 4-hydroxyphenylpyruvate | Amino Acid | 0.072181 | 0.979430302 |
| phenol sulfate | Amino Acid | 0.231738 | 1.049621673 |
| 4-hydroxyphenylacetate sulfate | Amino Acid | 0.399877 | 1.05341604 |
| N-formylphenylalanine | Amino Acid | 0.506325 | 1.002628415 |
| gentisate | Amino Acid | 0.572254 | 0.993821393 |
| tyrosine | Amino Acid | 0.631732 | 0.997896543 |
| vanillactate | Amino Acid | 0.70053 | 1.010672347 |
| N-acetyltyrosine | Amino Acid | 0.741123 | 0.991740254 |
| tyramine | Amino Acid | 0.79409 | 1.004962417 |
| 3-(4-hydroxyphenyl)lactate | Amino Acid | 0.924212 | 1.001461052 |
| trans-4-hydroxyproline | Amino Acid | 0.014716 | 0.987789428 |
| 2-oxoarginine* | Amino Acid | 0.147666 | 0.973489551 |
| N-acetylproline | Amino Acid | 0.221284 | 1.018224425 |
| N-acetylcitrulline | Amino Acid | 0.260042 | 0.980137605 |
| homocitrulline | Amino Acid | 0.281525 | 0.980066457 |
| dimethylarginine (SDMA + ADMA) | Amino Acid | 0.306881 | 1.010298206 |
| N-acetylarginine | Amino Acid | 0.514961 | 0.991322677 |

| Biochemical compound | Pathway | P-value (t-test) | Fold Change |
|---|---|---|---|
| citrulline | Amino Acid | 0.644351 | 0.997752846 |
| N-methylproline | Amino Acid | 0.748326 | 1.001134139 |
| N-alpha-acetylornithine | Amino Acid | 0.779908 | 0.995815723 |
| ornithine | Amino Acid | 0.802824 | 0.997511084 |
| N-delta-acetylornithine | Amino Acid | 0.889837 | 0.99773539 |
| arginine | Amino Acid | 0.894128 | 1.004322682 |
| argininate* | Amino Acid | 0.946499 | 1.001381187 |
| proline | Amino Acid | 0.948459 | 0.999805333 |
| argininosuccinate | Amino Acid | 0.997404 | 1.000301127 |
| N6-carboxymethyllysine | Carbohydrate | 0.449249 | 0.98806134 |
| N-acetylglucosaminylasparagine | Carbohydrate | 0.196288 | 1.023808075 |
| N-acetylglucosamine/N-acetylgalactosamine | Carbohydrate | 0.212093 | 1.021569532 |
| N-acetylmuramate | Carbohydrate | 0.274773 | 0.987984572 |
| N-acetylneuraminate | Carbohydrate | 0.307303 | 1.024437194 |
| glucuronate | Carbohydrate | 0.45345 | 1.014027512 |
| erythronate* | Carbohydrate | 0.671511 | 0.997368207 |
| N-acetyl-beta-glucosaminylamine | Carbohydrate | 0.957861 | 1.000454534 |
| raffinose | Carbohydrate | 0.509785 | 0.938998964 |
| sucrose | Carbohydrate | 0.629998 | 1.030943893 |
| mannitol/sorbitol | Carbohydrate | 0.328618 | 0.980373651 |
| fructose | Carbohydrate | 0.444513 | 0.973035433 |
| galactonate | Carbohydrate | 0.639911 | 1.015877838 |
| mannose | Carbohydrate | 0.659802 | 0.992536389 |
| galactitol (dulcitol) | Carbohydrate | 0.692566 | 0.993021623 |
| maltose | Carbohydrate | 0.853186 | 0.996711631 |
| 1,5-anhydroglucitol (1,5-AG) | Carbohydrate | 0.196122 | 1.016737741 |
| glucose 6-phosphate | Carbohydrate | 0.224251 | 1.086618564 |
| pyruvate | Carbohydrate | 0.283188 | 0.985168227 |
| lactate | Carbohydrate | 0.504335 | 0.992448693 |
| glucose | Carbohydrate | 0.726892 | 1.003246782 |
| glycerate | Carbohydrate | 0.911554 | 0.999332856 |
| arabonate/xylonate | Carbohydrate | 0.222075 | 0.979941501 |
| fucose | Carbohydrate | 0.23547 | 1.021464383 |
| ribose | Carbohydrate | 0.332837 | 0.99093598 |
| ribonate | Carbohydrate | 0.505424 | 1.013169922 |
| sedoheptulose | Carbohydrate | 0.522186 | 0.990457789 |
| xylose | Carbohydrate | 0.552386 | 1.011716957 |
| ribulonate/xylulonate* | Carbohydrate | 0.735063 | 1.004877308 |
| arabinose | Carbohydrate | 0.740684 | 1.0042687 |
| arabitol/xylitol | Carbohydrate | 0.843553 | 0.99549997 |

| Biochemical compound | Pathway | P-value (t-test) | Fold Change |
|---|---|---|---|
| ribulose/xylulose | Carbohydrate | 0.976648 | 1.00035779 |
| ribitol | Carbohydrate | 0.993902 | 1.000213252 |
| sedoheptulose-7-phosphate | Carbohydrate | 0.462573 | 1.053132138 |
| oxalate (ethanedioate) | Cofactors and Vitamins | 0.490282 | 0.992365366 |
| threonate | Cofactors and Vitamins | 0.542251 | 1.013107779 |
| biotin | Cofactors and Vitamins | 0.494027 | 0.98816979 |
| bilirubin (E,E)* | Cofactors and Vitamins | 0.029299 | 1.061286204 |
| bilirubin (Z,Z) | Cofactors and Vitamins | 0.074834 | 1.032367849 |
| protoporphyrin IX | Cofactors and Vitamins | 0.257851 | 0.980921294 |
| biliverdin | Cofactors and Vitamins | 0.331831 | 1.025134039 |
| D-urobilin | Cofactors and Vitamins | 0.371803 | 0.986106407 |
| I-urobilinogen | Cofactors and Vitamins | 0.376606 | 0.970148163 |
| heme | Cofactors and Vitamins | 0.576342 | 0.983637875 |
| L-urobilin | Cofactors and Vitamins | 0.808725 | 1.003992916 |
| N1-Methyl-2-pyridone-5-carboxamide | Cofactors and Vitamins | 0.111952 | 1.085218424 |
| nicotinamide riboside | Cofactors and Vitamins | 0.151538 | 1.029125644 |
| nicotinamide | Cofactors and Vitamins | 0.314731 | 0.988019863 |
| 6-hydroxynicotinate | Cofactors and Vitamins | 0.342226 | 0.96889412 |
| trigonelline (N'-methylnicotinate) | Cofactors and Vitamins | 0.398796 | 0.983132691 |
| nicotinate | Cofactors and Vitamins | 0.465686 | 0.996624399 |
| quinolinate | Cofactors and Vitamins | 0.610291 | 0.995963242 |
| nicotinate ribonucleoside | Cofactors and Vitamins | 0.795231 | 0.991770277 |
| N1-Methyl-4-pyridone-3-carboxamide | Cofactors and Vitamins | 0.961158 | 0.996299348 |
| pantethine | Cofactors and Vitamins | 0.102038 | 0.952975051 |
| pantetheine | Cofactors and Vitamins | 0.115945 | 0.980940326 |
| pantothenate | Cofactors and Vitamins | 0.587565 | 1.004534956 |
| riboflavin (Vitamin B2) | Cofactors and Vitamins | 0.299718 | 0.9902936 |
| flavin mononucleotide (FMN) | Cofactors and Vitamins | 0.631821 | 0.987152572 |
| flavin adenine dinucleotide (FAD) | Cofactors and Vitamins | 0.931339 | 1.000977043 |
| thiamin (Vitamin B1) | Cofactors and Vitamins | 0.321285 | 1.01104097 |
| hydroxymethylpyrimidine | Cofactors and Vitamins | 0.34404 | 0.98918588 |
| thiamin monophosphate | Cofactors and Vitamins | 0.741437 | 0.997303705 |
| gamma-tocopherol/beta-tocopherol | Cofactors and Vitamins | 0.016379 | 1.010246153 |
| alpha-tocotrienol | Cofactors and Vitamins | 0.040974 | 1.016694223 |
| delta-tocopherol | Cofactors and Vitamins | 0.085242 | 1.007403588 |
| alpha-tocopherol | Cofactors and Vitamins | 0.115662 | 1.005192412 |
| gamma-CEHC sulfate* | Cofactors and Vitamins | 0.215066 | 1.029091198 |
| alpha-tocopherol acetate | Cofactors and Vitamins | 0.337261 | 1.008490548 |
| gamma-tocotrienol | Cofactors and Vitamins | 0.386574 | 1.007938802 |
| gamma-CEHC | Cofactors and Vitamins | 0.978925 | 1.000682247 |

| Biochemical compound | Pathway | P-value (t-test) | Fold Change |
|---|---|---|---|
| beta-cryptoxanthin | Cofactors and Vitamins | 0.097022 | 1.010153545 |
| carotene diol (2) | Cofactors and Vitamins | 0.159504 | 1.007121836 |
| carotene diol (3) | Cofactors and Vitamins | 0.415428 | 1.003330908 |
| carotene diol (1) | Cofactors and Vitamins | 0.430775 | 1.004024191 |
| retinol (Vitamin A) | Cofactors and Vitamins | 0.799658 | 1.005401182 |
| pyridoxal | Cofactors and Vitamins | 0.35863 | 0.993198377 |
| pyridoxate | Cofactors and Vitamins | 0.358748 | 1.002497838 |
| pyridoxine (Vitamin B6) | Cofactors and Vitamins | 0.404944 | 1.012220995 |
| pyridoxamine | Cofactors and Vitamins | 0.587335 | 0.991274574 |
| phosphate | Energy | 0.379748 | 0.982269081 |
| alpha-ketoglutarate | Energy | 0.053662 | 0.977652833 |
| aconitate [cis or trans] | Energy | 0.075666 | 0.982471708 |
| tricarballylate | Energy | 0.36902 | 0.990548424 |
| succinate | Energy | 0.483834 | 0.993397864 |
| fumarate | Energy | 0.519489 | 1.011377473 |
| citrate | Energy | 0.642162 | 0.995775715 |
| malate | Energy | 0.734848 | 0.994875922 |
| 2-methylcitrate/homocitrate | Energy | 0.740355 | 0.996367696 |
| isocitric lactone | Energy | 0.980436 | 0.999081507 |
| deoxycarnitine | Lipid | 0.268836 | 0.983702108 |
| carnitine | Lipid | 0.902772 | 0.998240975 |
| ceramide (d18:1/20:0, d16:1/22:0, d20:1/18:0)* | Lipid | 0.087873 | 1.018203487 |
| N-stearoyl-sphingosine (d18:1/18:0)* | Lipid | 0.113739 | 1.026400798 |
| ceramide (d18:2/24:1, d18:1/24:2)* | Lipid | 0.246021 | 1.018067115 |
| N-palmitoyl-sphingosine (d18:1/16:0) | Lipid | 0.27307 | 1.011971491 |
| N-(2-hydroxypalmitoyl)-sphingosine (d18:1/16:0(2OH)) | Lipid | 0.365021 | 0.980202097 |
| ceramide (d18:1/17:0, d17:1/18:0)* | Lipid | 0.373652 | 1.023286635 |
| ceramide (d18:1/14:0, d16:1/16:0)* | Lipid | 0.966882 | 1.000670721 |
| linoleoyl-linoleoyl-glycerol (18:2/18:2) [2]* | Lipid | 0.171416 | 1.031871224 |
| oleoyl-linolenoyl-glycerol (18:1/18:3) [2]* | Lipid | 0.231657 | 1.023306954 |
| palmitoyl-myristoyl-glycerol (16:0/14:0) [2] | Lipid | 0.30913 | 1.008103088 |
| oleoyl-linoleoyl-glycerol (18:1/18:2) [1] | Lipid | 0.328043 | 1.018756816 |
| stearoyl-linoleoyl-glycerol (18:0/18:2) [1]* | Lipid | 0.332448 | 1.029553792 |
| linoleoyl-linolenoyl-glycerol (18:2/18:3) [1]* | Lipid | 0.393 | 1.018923564 |
| oleoyl-oleoyl-glycerol (18:1/18:1) [1]* | Lipid | 0.394693 | 1.020096441 |
| palmitoyl-linoleoyl-glycerol (16:0/18:2) [1]* | Lipid | 0.405225 | 1.016357365 |
| linoleoyl-linoleoyl-glycerol (18:2/18:2) [1]* | Lipid | 0.430851 | 1.017813228 |
| linoleoyl-arachidonoyl-glycerol (18:2/20:4) [1]* | Lipid | 0.47385 | 1.022357096 |

| Biochemical compound | Pathway | P-value (t-test) | Fold Change |
|---|---|---|---|
| palmitoyl-linolenoyl-glycerol (16:0/18:3) [2]* | Lipid | 0.478389 | 1.018849281 |
| oleoyl-oleoyl-glycerol (18:1/18:1) [2]* | Lipid | 0.517061 | 1.015414561 |
| diacylglycerol (16:1/18:2 [2], 16:0/18:3 [1])* | Lipid | 0.53454 | 1.017795476 |
| palmitoyl-linoleoyl-glycerol (16:0/18:2) [2]* | Lipid | 0.543773 | 1.01185428 |
| palmitoyl-palmitoyl-glycerol (16:0/16:0) [2]* | Lipid | 0.557948 | 1.009039273 |
| oleoyl-linoleoyl-glycerol (18:1/18:2) [2] | Lipid | 0.558268 | 1.010661947 |
| palmitoyl-oleoyl-glycerol (16:0/18:1) [1]* | Lipid | 0.565277 | 1.011127713 |
| linoleoyl-linolenoyl-glycerol (18:2/18:3) [2]* | Lipid | 0.640798 | 1.010767301 |
| palmitoyl-arachidonoyl-glycerol (16:0/20:4) [2]* | Lipid | 0.653645 | 1.012585631 |
| palmitoleoyl-linoleoyl-glycerol (16:1/18:2) [1]* | Lipid | 0.663321 | 1.017971567 |
| linoleoyl-arachidonoyl-glycerol (18:2/20:4) [2]* | Lipid | 0.700954 | 1.01954465 |
| linolenoyl-linolenoyl-glycerol (18:3/18:3) [2]* | Lipid | 0.713576 | 1.010576146 |
| palmitoyl-oleoyl-glycerol (16:0/18:1) [2]* | Lipid | 0.746956 | 1.005854096 |
| oleoyl-arachidonoyl-glycerol (18:1/20:4) [2]* | Lipid | 0.793142 | 1.017528673 |
| diacylglycerol (14:0/18:1, 16:0/16:1) [1]* | Lipid | 0.820341 | 0.993684992 |
| diacylglycerol (12:0/18:1, 14:0/16:1, 16:0/14:1) [2]* | Lipid | 0.854478 | 0.989425001 |
| stearoyl-linoleoyl-glycerol (18:0/18:2) [2]* | Lipid | 0.941283 | 0.998286384 |
| diacylglycerol (14:0/18:1, 16:0/16:1) [2]* | Lipid | 0.997481 | 1.000075275 |
| N-palmitoyl-sphinganine (d18:0/16:0) | Lipid | 0.443093 | 1.006183605 |
| N-stearoyl-sphinganine (d18:0/18:0)* | Lipid | 0.765586 | 1.006497557 |
| palmitoyl dihydrosphingomyelin (d18:0/16:0)* | Lipid | 0.375506 | 1.009786172 |
| linoleoyl ethanolamide | Lipid | 0.00372 | 1.029836368 |
| linolenoyl ethanolamide | Lipid | 0.0188 | 1.034021742 |
| oleoyl ethanolamide | Lipid | 0.07746 | 1.022830623 |
| lignoceroyl ethanolamide (24:0)* | Lipid | 0.109767 | 1.03762379 |
| stearoyl ethanolamide | Lipid | 0.150006 | 0.975246013 |
| N-linoleoyltaurine* | Lipid | 0.154774 | 1.043885742 |
| arachidoyl ethanolamide (20:0)* | Lipid | 0.226002 | 0.972403583 |
| N-stearoyltaurine | Lipid | 0.243061 | 0.959284484 |
| margaroyl ethanolamide* | Lipid | 0.299831 | 0.968096912 |
| N-oleoyltaurine | Lipid | 0.996276 | 1.000145054 |
| OAHSA (18:1/OH-18:0) | Lipid | 0.082695 | 1.031122319 |
| LAHSA (18:2/OH-18:0)* | Lipid | 0.119619 | 1.019157782 |
| linoleoylcholine* | Lipid | 0.070049 | 1.072857831 |
| methylmalonate (MMA) | Lipid | 0.776485 | 0.995893699 |
| butyrylcarnitine (C4) | Lipid | 0.98168 | 1.000515142 |

| Biochemical compound | Pathway | P-value (t-test) | Fold Change |
|---|---|---|---|
| stearoylcarnitine (C18) | Lipid | 0.22844 | 1.01795103 |
| myristoleoylcarnitine (C14:1)* | Lipid | 0.240842 | 0.968009594 |
| dihomo-linoleoylcarnitine (C20:2)* | Lipid | 0.395918 | 1.011401828 |
| acetylcarnitine (C2) | Lipid | 0.54164 | 1.015357135 |
| palmitoleoylcarnitine (C16:1)* | Lipid | 0.616172 | 0.987305353 |
| linoleoylcarnitine (C18:2)* | Lipid | 0.65208 | 1.014232809 |
| oleoylcarnitine (C18:1) | Lipid | 0.752489 | 1.005670663 |
| arachidonoylcarnitine (C20:4) | Lipid | 0.769983 | 1.007017534 |
| octadecanedioylcarnitine (C18-DC)* | Lipid | 0.906612 | 1.003241169 |
| octadecenedioylcarnitine (C18:1-DC)* | Lipid | 0.912247 | 1.00469255 |
| margaroylcarnitine (C17)* | Lipid | 0.927182 | 1.002048145 |
| palmitoylcarnitine (C16) | Lipid | 0.987587 | 0.999674329 |
| valerylglycine | Lipid | 0.297288 | 1.006695497 |
| N-linoleoylglycine | Lipid | 0.561927 | 1.025630006 |
| malonate | Lipid | 0.983846 | 1.00019435 |
| (14 or 15)-methylpalmitate (a17:0 or i17:0) | Lipid | 0.111258 | 0.979899778 |
| (16 or 17)-methylstearate (a19:0 or i19:0) | Lipid | 0.493476 | 0.988088873 |
| (12 or 13)-methylmyristate (a15:0 or i15:0) | Lipid | 0.923853 | 0.998602915 |
| suberate (C8-DC) | Lipid | 0.007619 | 0.987119582 |
| 2-hydroxyglutarate | Lipid | 0.054273 | 0.971359271 |
| pimelate (C7-DC) | Lipid | 0.075625 | 0.976873656 |
| glutarate (C5-DC) | Lipid | 0.142155 | 0.980219627 |
| azelate (C9-DC) | Lipid | 0.144072 | 0.983400674 |
| dodecanedioate (C12-DC) | Lipid | 0.168693 | 0.984189696 |
| octadecanedioate (C18-DC) | Lipid | 0.195826 | 0.984124669 |
| 2-hydroxyadipate | Lipid | 0.19785 | 0.975528762 |
| docosadioate (C22-DC) | Lipid | 0.269112 | 0.988634528 |
| sebacate (C10-DC) | Lipid | 0.269255 | 0.991152848 |
| adipate (C6-DC) | Lipid | 0.326179 | 0.98577766 |
| 3-methyladipate | Lipid | 0.44377 | 1.017393693 |
| 3-hydroxyadipate* | Lipid | 0.46328 | 0.989181347 |
| undecanedioate (C11-DC) | Lipid | 0.525279 | 0.994989124 |
| 3-methylglutarate/2-methylglutarate | Lipid | 0.546743 | 0.994625756 |
| hexadecenedioate (C16:1-DC)* | Lipid | 0.555105 | 0.994576876 |
| dimethylmalonic acid | Lipid | 0.692332 | 0.997021171 |
| octadecenedioate (C18:1-DC)* | Lipid | 0.692795 | 0.994587819 |
| octadecadienedioate (C18:2-DC)* | Lipid | 0.81498 | 1.004805948 |
| hexadecanedioate (C16-DC) | Lipid | 0.815983 | 0.997570067 |
| dodecenedioate (C12:1-DC)* | Lipid | 0.858733 | 1.001721179 |
| 12,13-DiHOME | Lipid | 0.435518 | 1.007780889 |

| Biochemical compound | Pathway | P-value (t-test) | Fold Change |
|---|---|---|---|
| 9,10-DiHOME | Lipid | 0.485259 | 1.009190952 |
| 10-hydroxystearate | Lipid | 0.104926 | 1.016823335 |
| 2-hydroxylignocerate* | Lipid | 0.263626 | 0.98739238 |
| 3-hydroxysebacate | Lipid | 0.280762 | 0.995477939 |
| 3-hydroxypalmitate | Lipid | 0.284307 | 0.988068875 |
| 5-hydroxyhexanoate | Lipid | 0.36845 | 1.018238877 |
| 16-hydroxypalmitate | Lipid | 0.377234 | 1.008378882 |
| 3-hydroxystearate | Lipid | 0.409297 | 0.989938254 |
| 13-HODE + 9-HODE | Lipid | 0.460693 | 1.00584747 |
| 2-hydroxypalmitate | Lipid | 0.497369 | 0.985937465 |
| 3-hydroxymyristate | Lipid | 0.505058 | 1.011500346 |
| 3-hydroxyhexanoate | Lipid | 0.555969 | 1.005479262 |
| 3-hydroxysuberate | Lipid | 0.557182 | 1.003121478 |
| 2-hydroxybehenate | Lipid | 0.568897 | 0.991817621 |
| 4-hydroxybutyrate (GHB) | Lipid | 0.582392 | 1.013611501 |
| 2-hydroxynervonate* | Lipid | 0.673413 | 0.992687338 |
| 2-hydroxystearate | Lipid | 0.729219 | 0.994901766 |
| 2-hydroxyarachidate* | Lipid | 0.754394 | 1.004697747 |
| 3-hydroxylaurate | Lipid | 0.776058 | 1.00206307 |
| 3-hydroxyoctanoate | Lipid | 0.77718 | 0.996242209 |
| 2-hydroxyheptanoate* | Lipid | 0.923287 | 1.00221725 |
| glycerol | Lipid | 0.081391 | 1.012221945 |
| glycerophosphoglycerol | Lipid | 0.543021 | 0.995630754 |
| glycerol 3-phosphate | Lipid | 0.731757 | 0.996518499 |
| 1,2-dilinoleoyl-digalactosylglycerol (18:2/18:2)* | Lipid | 0.011453 | 1.016990862 |
| 1,2-dilinoleoyl-galactosylglycerol (18:2/18:2)* | Lipid | 0.023423 | 1.023983049 |
| 1-linoleoyl-2-linolenoyl-galactosylglycerol (18:2/18:3)* | Lipid | 0.072459 | 1.028903899 |
| 2-palmitoyl-galactosylglycerol (16:0)* | Lipid | 0.13748 | 0.976836245 |
| 1-linoleoyl-2-linolenoyl-digalactosylglycerol (18:2/18:3)* | Lipid | 0.174591 | 1.025870622 |
| 1-palmitoyl-galactosylglycerol (16:0)* | Lipid | 0.27995 | 0.981079315 |
| 1-palmitoyl-2-linoleoyl-galactosylglycerol (16:0/18:2)* | Lipid | 0.809386 | 0.997976052 |
| 1-palmitoyl-2-linoleoyl-digalactosylglycerol (16:0/18:2)* | Lipid | 0.905683 | 0.998694189 |
| glycosyl-N-palmitoyl-sphingosine (d18:1/16:0) | Lipid | 0.278516 | 1.016093822 |
| inositol hexakisphosphate | Lipid | 0.71071 | 1.025109541 |

| Biochemical compound | Pathway | P-value (t-test) | Fold Change |
|---|---|---|---|
| chiro-inositol | Lipid | 0.803455 | 0.992752239 |
| myo-inositol | Lipid | 0.819884 | 0.99568143 |
| 3-hydroxybutyrate (BHBA) | Lipid | 0.969341 | 1.001666675 |
| heptadecatrienoate (17:3)* | Lipid | 0.017057 | 1.024691248 |
| stearate (18:0) | Lipid | 0.091531 | 0.987212288 |
| trans-nonadecenoate (tr 19:1)* | Lipid | 0.111517 | 0.969880069 |
| myristate (14:0) | Lipid | 0.118961 | 0.990757951 |
| palmitate (16:0) | Lipid | 0.139539 | 0.988547681 |
| pentadecanoate (15:0) | Lipid | 0.188073 | 0.981728144 |
| margarate (17:0) | Lipid | 0.215532 | 0.981332 |
| nonadecanoate (19:0) | Lipid | 0.233185 | 0.98168997 |
| arachidate (20:0) | Lipid | 0.250981 | 0.986111422 |
| 10-nonadecenoate (19:1n9) | Lipid | 0.461485 | 0.991145941 |
| oleate/vaccenate (18:1) | Lipid | 0.489381 | 0.995837444 |
| erucate (22:1n9) | Lipid | 0.50784 | 0.993161277 |
| 10-heptadecenoate (17:1n7) | Lipid | 0.564538 | 0.994152911 |
| palmitoleate (16:1n7) | Lipid | 0.740103 | 0.994726209 |
| myristoleate (14:1n5) | Lipid | 0.862619 | 0.997566242 |
| eicosenoate (20:1) | Lipid | 0.917396 | 1.001111404 |
| 1-lignoceroyl-GPC (24:0) | Lipid | 0.002615 | 1.056459907 |
| 1-oleoyl-GPC (18:1) | Lipid | 0.013467 | 1.034327387 |
| 1-linoleoyl-GPC (18:2) | Lipid | 0.016602 | 1.035658474 |
| 1-stearoyl-GPC (18:0) | Lipid | 0.10079 | 1.024060473 |
| 1-palmitoyl-GPC (16:0) | Lipid | 0.126789 | 1.019422848 |
| 1-stearoyl-GPS (18:0)* | Lipid | 0.158388 | 1.025820941 |
| 1-linoleoyl-GPG (18:2)* | Lipid | 0.173327 | 0.966995004 |
| 1-palmitoyl-GPE (16:0) | Lipid | 0.209805 | 1.023077249 |
| 1-oleoyl-GPE (18:1) | Lipid | 0.241327 | 1.027262871 |
| 1-stearoyl-GPE (18:0) | Lipid | 0.333253 | 1.017146146 |
| 1-stearoyl-GPA (18:0) | Lipid | 0.388414 | 0.98151966 |
| 1-palmitoyl-GPA (16:0) | Lipid | 0.442843 | 0.983418226 |
| 1-palmitoyl-GPS (16:0)* | Lipid | 0.44957 | 0.984952414 |
| 1-oleoyl-GPG (18:1)* | Lipid | 0.525788 | 0.986246864 |
| 1-oleoyl-GPA (18:1) | Lipid | 0.525871 | 0.978815181 |
| 1-linoleoyl-GPA (18:2)* | Lipid | 0.539013 | 0.985766297 |
| 2-stearoyl-GPE (18:0)* | Lipid | 0.608821 | 0.980852305 |
| 1-stearoyl-GPI (18:0) | Lipid | 0.672609 | 0.988985377 |
| 1-palmitoyl-GPG (16:0)* | Lipid | 0.74946 | 0.997600348 |
| 1-stearoyl-GPG (18:0) | Lipid | 0.815628 | 0.995281267 |
| 1-linoleoyl-GPE (18:2)* | Lipid | 0.932258 | 0.998377985 |

| Biochemical compound | Pathway | P-value (t-test) | Fold Change |
|---|---|---|---|
| 1-palmitoyl-GPI (16:0) | Lipid | 0.962572 | 1.001226414 |
| 1-(1-enyl-stearoyl)-GPE (P-18:0)* | Lipid | 0.408981 | 1.015603376 |
| 1-(1-enyl-palmitoyl)-GPE (P-16:0)* | Lipid | 0.423232 | 0.975542466 |
| 1-(1-enyl-oleoyl)-GPE (P-18:1)* | Lipid | 0.705968 | 0.983807257 |
| caprate (10:0) | Lipid | 0.057281 | 0.988595081 |
| 3-hydroxy-3-methylglutarate | Lipid | 0.544242 | 0.992731045 |
| mevalonate | Lipid | 0.67977 | 0.99516335 |
| mevalonolactone | Lipid | 0.831632 | 0.99548213 |
| 1-linoleoylglycerol (18:2) | Lipid | 0.160409 | 1.014217785 |
| 1-eicosenoylglycerol (20:1) | Lipid | 0.170893 | 1.015320732 |
| 1-heptadecenoylglycerol (17:1)* | Lipid | 0.228741 | 0.976242653 |
| 1-stearoylglycerol (18:0) | Lipid | 0.239873 | 1.010979807 |
| 2-myristoylglycerol (14:0) | Lipid | 0.244602 | 0.974452207 |
| 1-oleoylglycerol (18:1) | Lipid | 0.257541 | 1.008899037 |
| 1-linolenoylglycerol (18:3) | Lipid | 0.258698 | 1.010327985 |
| 2-linoleoylglycerol (18:2) | Lipid | 0.263982 | 1.008444838 |
| 1-myristoylglycerol (14:0) | Lipid | 0.347618 | 0.985369303 |
| 2-palmitoleoylglycerol (16:1)* | Lipid | 0.373473 | 0.969968254 |
| 2-stearoylglycerol (18:0) | Lipid | 0.442731 | 0.993183079 |
| 2-palmitoylglycerol (16:0) | Lipid | 0.470709 | 1.006517005 |
| 1-dihomo-linolenylglycerol (20:3) | Lipid | 0.527323 | 1.016097965 |
| 1-palmitoylglycerol (16:0) | Lipid | 0.655102 | 1.00362925 |
| 1-margaroylglycerol (17:0) | Lipid | 0.726478 | 1.006014645 |
| 1-dihomo-linoleoylglycerol (20:2) | Lipid | 0.81577 | 1.003187541 |
| 1-docosahexaenoylglycerol (22:6) | Lipid | 0.867847 | 1.006650041 |
| 1-arachidonylglycerol (20:4) | Lipid | 0.879146 | 1.003286878 |
| 1-palmitoleoylglycerol (16:1)* | Lipid | 0.903429 | 1.002551474 |
| 2-arachidonoylglycerol (20:4) | Lipid | 0.943116 | 1.002837894 |
| 1-pentadecanoylglycerol (15:0) | Lipid | 0.948984 | 0.998655775 |
| 2-oleoylglycerol (18:1) | Lipid | 0.982915 | 1.000212721 |
| 1-palmitoyl-2-docosahexaenoyl-GPC (16:0/22:6) | Lipid | 0.069022 | 1.030222208 |
| 1-palmitoyl-2-stearoyl-GPC (16:0/18:0) | Lipid | 0.090858 | 1.049084097 |
| 1-oleoyl-2-linoleoyl-GPC (18:1/18:2)* | Lipid | 0.129363 | 1.025167889 |
| 1-myristoyl-2-palmitoyl-GPC (14:0/16:0) | Lipid | 0.134609 | 1.033897274 |
| 1,2-dipalmitoyl-GPC (16:0/16:0) | Lipid | 0.139439 | 1.026200207 |
| 1,2-dilinoleoyl-GPC (18:2/18:2) | Lipid | 0.178809 | 1.02580337 |
| 1-palmitoyl-2-linoleoyl-GPC (16:0/18:2) | Lipid | 0.242276 | 1.016704515 |
| 1-linoleoyl-2-arachidonoyl-GPC (18:2/20:4n6)* | Lipid | 0.567567 | 1.037572931 |

| Biochemical compound | Pathway | P-value (t-test) | Fold Change |
|---|---|---|---|
| 1-palmitoyl-2-arachidonoyl-GPC (16:0/20:4n6) | Lipid | 0.590561 | 1.01185114 |
| 1-linoleoyl-2-linolenoyl-GPC (18:2/18:3)* | Lipid | 0.598022 | 1.010351173 |
| 1-palmitoyl-2-oleoyl-GPC (16:0/18:1) | Lipid | 0.741297 | 1.005556979 |
| 1-stearoyl-2-arachidonoyl-GPC (18:0/20:4) | Lipid | 0.75494 | 1.011019624 |
| 1-palmitoyl-2-palmitoleoyl-GPC (16:0/16:1)* | Lipid | 0.843833 | 0.994947531 |
| 1-palmitoyl-2-dihomo-linolenoyl-GPC (16:0/20:3n3 or 6)* | Lipid | 0.905351 | 1.002940654 |
| 1-stearoyl-2-oleoyl-GPC (18:0/18:1) | Lipid | 0.942524 | 1.001858094 |
| 1-oleoyl-2-linoleoyl-GPE (18:1/18:2)* | Lipid | 0.055876 | 1.04092198 |
| 1-palmitoyl-2-linoleoyl-GPE (16:0/18:2) | Lipid | 0.055944 | 1.029497314 |
| 1-stearoyl-2-linoleoyl-GPE (18:0/18:2)* | Lipid | 0.6198 | 0.97833262 |
| 1-palmitoyl-2-arachidonoyl-GPE (16:0/20:4)* | Lipid | 0.685787 | 1.007416376 |
| 1-stearoyl-2-arachidonoyl-GPE (18:0/20:4) | Lipid | 0.707789 | 1.011824901 |
| 1,2-dilinoleoyl-GPE (18:2/18:2)* | Lipid | 0.866615 | 1.006011838 |
| 1-palmitoyl-2-oleoyl-GPE (16:0/18:1) | Lipid | 0.936281 | 0.997799448 |
| 1-palmitoyl-2-docosahexaenoyl-GPE (16:0/22:6)* | Lipid | 0.943377 | 0.99806766 |
| 1,2-dipalmitoyl-GPG (16:0/16:0) | Lipid | 0.522242 | 1.011463899 |
| 1-palmitoyl-2-linoleoyl-GPI (16:0/18:2) | Lipid | 0.426183 | 0.984610128 |
| 1-stearoyl-2-arachidonoyl-GPI (18:0/20:4) | Lipid | 0.755438 | 0.981833777 |
| 1-stearoyl-2-oleoyl-GPS (18:0/18:1) | Lipid | 0.904247 | 1.00195495 |
| glycerophosphoserine* | Lipid | 0.137706 | 1.02879105 |
| trimethylamine N-oxide | Lipid | 0.310095 | 1.018365448 |
| glycerophosphoethanolamine | Lipid | 0.370423 | 1.014676839 |
| choline | Lipid | 0.436777 | 1.005164113 |
| glycerophosphoinositol* | Lipid | 0.629328 | 1.009582487 |
| choline phosphate | Lipid | 0.930804 | 1.003176963 |
| glycerophosphorylcholine (GPC) | Lipid | 0.977175 | 0.99850426 |
| 1-(1-enyl-palmitoyl)-2-linoleoyl-GPC (P-16:0/18:2)* | Lipid | 0.335026 | 1.038977985 |
| 1-(1-enyl-palmitoyl)-2-arachidonoyl-GPC (P-16:0/20:4)* | Lipid | 0.432379 | 1.028281107 |
| 1-(1-enyl-palmitoyl)-2-linoleoyl-GPE (P-16:0/18:2)* | Lipid | 0.600501 | 1.027206805 |
| 1-(1-enyl-palmitoyl)-2-oleoyl-GPE (P-16:0/18:1)* | Lipid | 0.765402 | 0.995776359 |
| 1-(1-enyl-stearoyl)-2-arachidonoyl-GPE (P-18:0/20:4)* | Lipid | 0.814582 | 0.992915693 |
| 1-(1-enyl-palmitoyl)-2-oleoyl-GPC (P-16:0/18:1)* | Lipid | 0.837521 | 1.0059698 |

| Biochemical compound | Pathway | P-value (t-test) | Fold Change |
|---|---|---|---|
| 1-(1-enyl-palmitoyl)-2-arachidonoyl-GPE (P-16:0/20:4)* | Lipid | 0.919228 | 0.996849776 |
| linolenate [alpha or gamma; (18:3n3 or 6)] | Lipid | 0.050741 | 1.013845891 |
| docosapentaenoate (n6 DPA; 22:5n6) | Lipid | 0.12843 | 0.972819317 |
| stearidonate (18:4n3) | Lipid | 0.214196 | 1.063912904 |
| linoleate (18:2n6) | Lipid | 0.371857 | 1.004894343 |
| hexadecadienoate (16:2n6) | Lipid | 0.385708 | 1.013050051 |
| dihomo-linoleate (20:2n6) | Lipid | 0.459755 | 0.990855397 |
| mead acid (20:3n9) | Lipid | 0.480161 | 0.985943799 |
| adrenate (22:4n6) | Lipid | 0.520175 | 1.008606204 |
| docosadienoate (22:2n6) | Lipid | 0.62353 | 0.991347145 |
| hexadecatrienoate (16:3n3) | Lipid | 0.713208 | 1.011388039 |
| dihomo-linolenate (20:3n3 or n6) | Lipid | 0.73427 | 0.995414583 |
| eicosapentaenoate (EPA; 20:5n3) | Lipid | 0.756418 | 1.003928774 |
| docosatrienoate (22:3n6)* | Lipid | 0.816414 | 0.993615384 |
| nisinate (24:6n3) | Lipid | 0.861514 | 0.995567314 |
| arachidonate (20:4n6) | Lipid | 0.888877 | 0.998431717 |
| docosahexaenoate (DHA; 22:6n3) | Lipid | 0.905175 | 1.001739225 |
| docosapentaenoate (n3 DPA; 22:5n3) | Lipid | 0.997362 | 0.999940602 |
| pregnenediol disulfate (C21H34O8S2)* | Lipid | 0.787375 | 1.008934268 |
| glycocholate | Lipid | 0.165063 | 1.037154963 |
| tauro-beta-muricholate | Lipid | 0.188153 | 1.032453248 |
| chenodeoxycholate | Lipid | 0.228179 | 1.025745771 |
| glyco-beta-muricholate** | Lipid | 0.402444 | 1.020180689 |
| cholate | Lipid | 0.410146 | 1.023010428 |
| alpha-muricholate | Lipid | 0.474784 | 1.014410457 |
| beta-muricholate | Lipid | 0.537454 | 1.007960063 |
| taurocholate | Lipid | 0.878448 | 1.004826896 |
| taurochenodeoxycholate | Lipid | 0.975732 | 1.000684919 |
| 5alpha-pregnan-3beta,20alpha-diol disulfate | Lipid | 0.724278 | 1.014117383 |
| 6-beta-hydroxylithocholate | Lipid | 0.061595 | 1.024834575 |
| taurohyodeoxycholic acid | Lipid | 0.069142 | 1.062604192 |
| isohyodeoxycholate | Lipid | 0.225539 | 1.027804969 |
| dehydrolithocholate | Lipid | 0.229213 | 0.974108001 |
| hyocholate | Lipid | 0.230025 | 1.038321987 |
| ursodeoxycholate | Lipid | 0.238346 | 1.028738288 |
| tauroursodeoxycholate | Lipid | 0.299465 | 1.029349985 |
| deoxycholic acid sulfate | Lipid | 0.433298 | 0.973489597 |
| 7-ketodeoxycholate | Lipid | 0.568991 | 1.011087459 |
| lithocholate | Lipid | 0.622486 | 0.992349456 |

| Biochemical compound | Pathway | P-value (t-test) | Fold Change |
|---|---|---|---|
| 3b-hydroxy-5-cholenoic acid | Lipid | 0.655723 | 1.015393612 |
| 7,12-diketolithocholate | Lipid | 0.733342 | 1.014331784 |
| deoxycholate | Lipid | 0.746646 | 1.006065861 |
| 6-oxolithocholate | Lipid | 0.765019 | 1.006834193 |
| taurohyocholate* | Lipid | 0.773008 | 1.020281151 |
| 12-dehydrocholate | Lipid | 0.773814 | 1.012184084 |
| taurodeoxycholate | Lipid | 0.796963 | 1.010931664 |
| taurolithocholate 3-sulfate | Lipid | 0.822514 | 0.989808742 |
| taurochenodeoxycholate sulfate | Lipid | 0.83603 | 1.01182476 |
| taurocholenate sulfate* | Lipid | 0.84976 | 1.006827824 |
| 3-dehydrocholate | Lipid | 0.931566 | 0.996472392 |
| ursocholate | Lipid | 0.96271 | 0.999288896 |
| valerate (5:0) | Lipid | 0.969008 | 0.999545489 |
| phytosphingosine | Lipid | 0.046042 | 1.019928793 |
| 3-ketosphinganine | Lipid | 0.377353 | 1.016152739 |
| sphingadienine | Lipid | 0.730915 | 0.995408351 |
| sphinganine | Lipid | 0.82533 | 1.003365915 |
| palmitoyl sphingomyelin (d18:1/16:0) | Lipid | 0.264205 | 1.01281314 |
| stearoyl sphingomyelin (d18:1/18:0) | Lipid | 0.362268 | 1.027368392 |
| sphingomyelin (d17:1/16:0, d18:1/15:0, d16:1/17:0)* | Lipid | 0.363278 | 1.018636776 |
| sphingomyelin (d18:1/20:0, d16:1/22:0)* | Lipid | 0.652694 | 1.032602583 |
| hexadecasphingosine (d16:1)* | Lipid | 0.335879 | 1.024771081 |
| eicosanoylsphingosine (d20:1)* | Lipid | 0.526127 | 1.007485108 |
| sphingosine | Lipid | 0.889308 | 0.997586035 |
| heptadecasphingosine (d17:1) | Lipid | 0.902849 | 1.002841386 |
| beta-sitosterol | Lipid | 0.01369 | 1.014459847 |
| stigmasterol | Lipid | 0.030937 | 1.012926246 |
| campesterol | Lipid | 0.038772 | 1.017016776 |
| cholesterol | Lipid | 0.042509 | 1.02118387 |
| ergosterol | Lipid | 0.35164 | 1.00953743 |
| 7-alpha-hydroxy-3-oxo-4-cholestenoate (7-Hoca) | Lipid | 0.364425 | 1.015537138 |
| 4-cholesten-3-one | Lipid | 0.691 | 1.002442804 |
| coprostanol | Lipid | 0.867053 | 0.995873191 |
| 3beta-hydroxy-5-cholestenoate | Lipid | 0.92733 | 0.997513902 |
| (3'-5')-cytidylyladenosine | Nucleotide | 0.253221 | 0.975930905 |
| (3'-5')-cytidylyluridine* | Nucleotide | 0.360656 | 0.988153467 |
| (3'-5')-uridylyluridine | Nucleotide | 0.410919 | 0.988052462 |
| (3'-5')-uridylylguanosine | Nucleotide | 0.550673 | 0.986109211 |

| Biochemical compound | Pathway | P-value (t-test) | Fold Change |
|---|---|---|---|
| (3'-5')-uridylylcytidine* | Nucleotide | 0.563026 | 0.989685449 |
| (3'-5')-cytidylylguanosine | Nucleotide | 0.612017 | 0.989520775 |
| (3'-5')-uridylyladenosine | Nucleotide | 0.633924 | 0.989768931 |
| (3'-5')-cytidylylcytidine* | Nucleotide | 0.641768 | 0.993118652 |
| (3'-5')-adenylylcytidine | Nucleotide | 0.873694 | 0.996491723 |
| (3'-5')-adenylyladenosine* | Nucleotide | 0.9384 | 0.998600428 |
| (3'-5')-guanylylcytidine | Nucleotide | 0.941689 | 0.997108444 |
| (3'-5')-guanylyluridine | Nucleotide | 0.95915 | 0.997752798 |
| (3'-5')-adenylyluridine | Nucleotide | 0.987141 | 0.999651782 |
| methylphosphate | Nucleotide | 0.739477 | 0.993061218 |
| 2'-deoxyinosine | Nucleotide | 0.156816 | 1.014183473 |
| urate | Nucleotide | 0.227064 | 1.017772949 |
| inosine | Nucleotide | 0.595924 | 1.004331185 |
| hypoxanthine | Nucleotide | 0.620016 | 0.998832871 |
| xanthine | Nucleotide | 0.758265 | 1.002136144 |
| xanthosine | Nucleotide | 0.770554 | 0.99537991 |
| allantoin | Nucleotide | 0.987082 | 1.000900783 |
| 2'-deoxyadenosine | Nucleotide | 0.043641 | 0.923940457 |
| N6-carbamoylthreonyladenosine | Nucleotide | 0.06527 | 1.022264978 |
| adenosine | Nucleotide | 0.072912 | 0.962690421 |
| adenine | Nucleotide | 0.102785 | 0.961134768 |
| 1-methyladenine | Nucleotide | 0.530339 | 1.00558537 |
| 2'-deoxyadenosine 5'-monophosphate | Nucleotide | 0.634272 | 1.010527994 |
| adenosine 5'-monophosphate (AMP) | Nucleotide | 0.826081 | 0.996985356 |
| N6-methyladenosine | Nucleotide | 0.840361 | 1.006569993 |
| N1-methyladenosine | Nucleotide | 0.863088 | 1.004473581 |
| N6-succinyladenosine | Nucleotide | 0.901423 | 0.996397077 |
| 2'-deoxyguanosine | Nucleotide | 0.183041 | 1.009932765 |
| N1-methylguanosine | Nucleotide | 0.351704 | 1.006354917 |
| guanine | Nucleotide | 0.442162 | 0.994756578 |
| guanosine-2',3'-cyclic monophosphate | Nucleotide | 0.508729 | 1.018714698 |
| guanosine 3'-monophosphate (3'-GMP) | Nucleotide | 0.522306 | 1.029897623 |
| guanosine | Nucleotide | 0.602666 | 0.99350526 |
| 1-methylguanine | Nucleotide | 0.927356 | 1.001566861 |
| 7-methylguanine | Nucleotide | 0.997628 | 0.999963018 |
| 5-methyl-2'-deoxycytidine | Nucleotide | 0.089439 | 1.02624951 |
| cytosine | Nucleotide | 0.193237 | 0.98650952 |
| 2'-deoxycytidine | Nucleotide | 0.303789 | 1.012292832 |
| cytidine 5'-monophosphate (5'-CMP) | Nucleotide | 0.44356 | 0.988621569 |
| 5-methylcytosine | Nucleotide | 0.500577 | 1.010845997 |

| Biochemical compound | Pathway | P-value (t-test) | Fold Change |
|---|---|---|---|
| 2'-deoxycytidine 5'-monophosphate | Nucleotide | 0.50313 | 1.011712668 |
| cytidine | Nucleotide | 0.916745 | 0.998947429 |
| N-carbamoylaspartate | Nucleotide | 0.258518 | 0.97985326 |
| orotate | Nucleotide | 0.520508 | 0.987374513 |
| dihydroorotate | Nucleotide | 0.538267 | 0.974056621 |
| thymidine | Nucleotide | 0.129042 | 1.015018074 |
| thymine | Nucleotide | 0.336851 | 1.012577312 |
| thymidine 5'-monophosphate | Nucleotide | 0.725093 | 1.010673795 |
| 2'-deoxyuridine | Nucleotide | 0.103955 | 1.015507527 |
| 5-methyluridine (ribothymidine) | Nucleotide | 0.146533 | 1.011021863 |
| uridine 5'-monophosphate (UMP) | Nucleotide | 0.340462 | 1.011145603 |
| 2'-O-methyluridine | Nucleotide | 0.35666 | 1.01614565 |
| 3-ureidoisobutyrate | Nucleotide | 0.461813 | 1.068978222 |
| 3-ureidopropionate | Nucleotide | 0.479052 | 1.017278654 |
| uridine | Nucleotide | 0.616658 | 1.00398568 |
| N-acetyl-beta-alanine | Nucleotide | 0.716866 | 1.011928411 |
| beta-alanine | Nucleotide | 0.76724 | 1.001773621 |
| uridine-2',3'-cyclic monophosphate | Nucleotide | 0.778889 | 0.987849749 |
| uracil | Nucleotide | 0.806817 | 1.001438831 |
| pseudouridine | Nucleotide | 0.963134 | 1.000496691 |
| 5,6-dihydrouridine | Nucleotide | 0.991497 | 1.000082466 |
| leucylglycine | Peptide | 0.278329 | 0.987373209 |
| valylglycine | Peptide | 0.362731 | 0.992764437 |
| prolylglycine | Peptide | 0.39579 | 1.040702072 |
| leucylglutamine* | Peptide | 0.445068 | 0.986882535 |
| valylglutamine | Peptide | 0.517674 | 0.989337057 |
| alanylleucine | Peptide | 0.675986 | 0.993856834 |
| leucylalanine | Peptide | 0.696973 | 0.990955034 |
| glycylleucine | Peptide | 0.70728 | 1.002191886 |
| tryptophylglycine | Peptide | 0.723464 | 1.014841901 |
| glutaminylleucine | Peptide | 0.740798 | 1.007941854 |
| phenylalanylglycine | Peptide | 0.758884 | 0.991509031 |
| isoleucylglycine | Peptide | 0.762617 | 1.004103669 |
| threonylphenylalanine | Peptide | 0.763086 | 0.996129659 |
| phenylalanylalanine | Peptide | 0.768356 | 0.993244659 |
| glycylvaline | Peptide | 0.775702 | 1.001385703 |
| tyrosylglycine | Peptide | 0.807157 | 0.991446871 |
| glycylisoleucine | Peptide | 0.823223 | 1.001609328 |
| lysylleucine | Peptide | 0.846558 | 1.004008417 |
| valylleucine | Peptide | 0.929077 | 0.99888692 |

| Biochemical compound | Pathway | P-value (t-test) | Fold Change |
|---|---|---|---|
| gamma-glutamyl-epsilon-lysine | Peptide | 0.045231 | 1.03936094 |
| gamma-glutamylglutamine | Peptide | 0.161721 | 0.982480364 |
| gamma-glutamylhistidine | Peptide | 0.264087 | 0.972268302 |
| gamma-glutamylmethionine | Peptide | 0.280204 | 0.987080005 |
| gamma-glutamylalanine | Peptide | 0.334544 | 0.989437568 |
| gamma-glutamyl-alpha-lysine | Peptide | 0.433562 | 0.989975553 |
| gamma-glutamylthreonine | Peptide | 0.521763 | 0.991258444 |
| gamma-glutamyltyrosine | Peptide | 0.535391 | 0.993166787 |
| gamma-glutamylleucine | Peptide | 0.549694 | 0.992150245 |
| gamma-glutamylglycine | Peptide | 0.611773 | 1.013054789 |
| gamma-glutamylphenylalanine | Peptide | 0.659759 | 0.994510309 |
| gamma-glutamylisoleucine* | Peptide | 0.6773 | 0.994525654 |
| gamma-glutamylglutamate | Peptide | 0.681205 | 0.992162321 |
| alanyl-glutamyl-meso-diaminopimelate | Peptide | 0.927198 | 0.998455056 |
| tartronate (hydroxymalonate) | Xenobiotics | 0.00236 | 0.948908087 |
| 3-deoxyoctulosonate | Xenobiotics | 0.258755 | 1.017327223 |
| 4-hydroxyhippurate | Xenobiotics | 0.046661 | 1.076585287 |
| catechol sulfate | Xenobiotics | 0.121791 | 1.07320693 |
| benzoate | Xenobiotics | 0.147425 | 0.978279273 |
| 3-(3-hydroxyphenyl)propionate sulfate | Xenobiotics | 0.42914 | 1.036138772 |
| 4-hydroxymandelate | Xenobiotics | 0.443421 | 0.99076818 |
| 3-phenylpropionate (hydrocinnamate) | Xenobiotics | 0.622819 | 0.995267556 |
| 3-(4-hydroxyphenyl)propionate | Xenobiotics | 0.713395 | 1.00384311 |
| hippurate | Xenobiotics | 0.847138 | 0.987756892 |
| 3-(3-hydroxyphenyl)propionate | Xenobiotics | 0.865591 | 0.995631984 |
| 2,4,6-trihydroxybenzoate | Xenobiotics | 0.952434 | 0.99937763 |
| 4-hydroxybenzoate | Xenobiotics | 0.982924 | 0.999489942 |
| 2-aminophenol sulfate | Xenobiotics | 0.176081 | 1.042080865 |
| succinimide | Xenobiotics | 0.209929 | 0.992188999 |
| thioproline | Xenobiotics | 0.450129 | 0.974957998 |
| 1,2,3-benzenetriol sulfate (1) | Xenobiotics | 0.466534 | 1.056370041 |
| S-(3-hydroxypropyl)mercapturic acid (HPMA) | Xenobiotics | 0.53221 | 0.989007468 |
| O-sulfo-L-tyrosine | Xenobiotics | 0.636928 | 0.982096437 |
| 4-thiouracil | Xenobiotics | 0.808551 | 0.997968565 |
| sulfate* | Xenobiotics | 0.913696 | 1.000781287 |
| 1,2,3-benzenetriol sulfate (2) | Xenobiotics | #DIV/0! | 1.008546866 |
| hydroquinone sulfate | Xenobiotics | 0.193962 | 1.030888636 |
| salicylate | Xenobiotics | 0.233388 | 0.986827355 |
| formononetin | Xenobiotics | 0.051165 | 1.144604759 |
| chrysoeriol (3'-O-methylluteolin) | Xenobiotics | 0.051436 | 1.094236916 |

| Biochemical compound | Pathway | P-value (t-test) | Fold Change |
|---|---|---|---|
| pyrraline | Xenobiotics | 0.065814 | 1.035381794 |
| pheophytin A | Xenobiotics | 0.090253 | 1.025212846 |
| gluconate | Xenobiotics | 0.121089 | 0.974286929 |
| stachydrine | Xenobiotics | 0.121752 | 1.027789725 |
| 2,8-quinolinediol sulfate | Xenobiotics | 0.138881 | 1.060445773 |
| apigenin | Xenobiotics | 0.192052 | 1.054511668 |
| genistein sulfate* | Xenobiotics | 0.195491 | 1.059260362 |
| 2-isopropylmalate | Xenobiotics | 0.200627 | 0.980020426 |
| enterodiol | Xenobiotics | 0.234504 | 1.020743248 |
| syringic acid | Xenobiotics | 0.25747 | 1.019828059 |
| ferulate | Xenobiotics | 0.27039 | 1.013851249 |
| ferulic acid 4-sulfate | Xenobiotics | 0.280148 | 1.056839191 |
| tyrosol | Xenobiotics | 0.281822 | 0.975876273 |
| caffeate | Xenobiotics | 0.294235 | 1.01947537 |
| daidzein sulfate (2) | Xenobiotics | 0.326897 | 1.122704986 |
| 2-keto-3-deoxy-gluconate | Xenobiotics | 0.337449 | 0.987556509 |
| naringenin | Xenobiotics | 0.384036 | 1.027795475 |
| genistein | Xenobiotics | 0.385264 | 1.017058989 |
| 3-dehydroshikimate | Xenobiotics | 0.41904 | 1.020212309 |
| indoleacrylate | Xenobiotics | 0.452551 | 1.017390595 |
| equol | Xenobiotics | 0.467735 | 0.973429126 |
| soyasaponin II | Xenobiotics | 0.521472 | 1.032476304 |
| soyasaponin I | Xenobiotics | 0.558914 | 1.020393033 |
| 1,1-kestotetraose | Xenobiotics | 0.577112 | 0.955047925 |
| equol sulfate | Xenobiotics | 0.590122 | 1.020420755 |
| quinate | Xenobiotics | 0.596772 | 0.99050761 |
| daidzein sulfate (1) | Xenobiotics | 0.606762 | 1.067622857 |
| soyasaponin III | Xenobiotics | 0.611394 | 1.031766 |
| N-glycolylneuraminate | Xenobiotics | 0.623552 | 0.976386817 |
| biochanin A | Xenobiotics | 0.632908 | 1.02171998 |
| daidzein | Xenobiotics | 0.641371 | 1.016698143 |
| indolin-2-one | Xenobiotics | 0.650798 | 1.015054685 |
| 2,8-quinolinediol | Xenobiotics | 0.690457 | 1.004313015 |
| dihydrocaffeate sulfate (2) | Xenobiotics | 0.722441 | 1.021214863 |
| sinapate | Xenobiotics | 0.728835 | 1.006325841 |
| ergothioneine | Xenobiotics | 0.773197 | 1.004578565 |
| glycitein | Xenobiotics | 0.787416 | 0.990114236 |
| 1H-quinolin-2-one | Xenobiotics | 0.827613 | 1.003891573 |
| dipicolinate | Xenobiotics | 0.830541 | 0.991673385 |
| histidine betaine (hercynine)* | Xenobiotics | 0.833395 | 0.991073976 |

| Biochemical compound | Pathway | P-value (t-test) | Fold Change |
|---|---|---|---|
| dihydroferulic acid | Xenobiotics | 0.858065 | 0.996987277 |
| 4-hydroxycinnamate | Xenobiotics | 0.866881 | 0.998707787 |
| nicotianamine | Xenobiotics | 0.889594 | 0.995913741 |
| 3,5-dihydroxybenzoic acid | Xenobiotics | 0.897692 | 1.009120323 |
| 2-oxindole-3-acetate | Xenobiotics | 0.90812 | 0.998356627 |
| apigenin 7-O(6-malonyl-beta-D-glucoside) | Xenobiotics | 0.922872 | 0.997588025 |
| 2,3-dihydroxyisovalerate | Xenobiotics | 0.926625 | 0.998991224 |
| enterolactone | Xenobiotics | 0.936031 | 0.998138971 |
| vanillate | Xenobiotics | 0.950789 | 1.001170954 |
| diaminopimelate | Xenobiotics | 0.959238 | 0.99893317 |
| 1,2-dilinolenoyl-digalactosylglycerol (18:3/18:3) | Xenobiotics | 0.995931 | 0.999863972 |
